# Aerodynamic characteristics of a feathered dinosaur measured using physical models. Effects of form on static stability and control effectiveness

**DOI:** 10.1101/001297

**Authors:** Dennis Evangelista, Griselda Cardona, Eric Guenther-Gleason, Tony Huynh, Austin Kwong, Dylan Marks, Neil Ray, Adrian Tisbe, Kyle Tse, Mimi Koehl

**Affiliations:** Department of Integrative Biology, University of California, Berkeley, CA, USA; Department of Mechanical Engineering, University of California, Berkeley, CA, USA; Department of Bioengineering, University of California, Berkeley, CA, USA

## Abstract

We report the effects of posture and morphology on the static aerodynamic stability and control effectiveness of physical models based on the feathered dinosaur, †*Microraptor gui*, from the Cretaceous of China. Postures had similar lift and drag coefficients and were broadly similar when simplified metrics of gliding were considered, but they exhibited different stability characteristics depending on the position of the legs and the presence of feathers on the legs and the tail. Both stability and the function of appendages in generating maneuvering forces and torques changed as the glide angle or angle of attack were changed. These are significant because they represent an aerial environment that may have shifted during the evolution of directed aerial descent and other aerial behaviors. Certain movements were particularly effective (symmetric movements of the wings and tail in pitch, asymmetric wing movements, some tail movements). Other appendages altered their function from creating yaws at high angle of attack to rolls at low angle of attack, or reversed their function entirely. While †*M. gui* lived after †*Archaeopteryx* and likely represents a side experiment with feathered morphology, the general patterns of stability and control effectiveness suggested from the manipulations of forelimb, hindlimb and tail morphology here may help understand the evolution of flight control aerodynamics in vertebrates. Though these results rest on a single specimen, as further fossils with different morphologies tested, the findings here could be applied in a phylogenetic context to reveal biomechanical constraints on extinct flyers arising from the need to maneuver.

## Introduction

The evolution of flight in vertebrates, and particularly in birds, is the subject of lively debate and considerable speculation. Furthermore, flight ability of extinct vertebrates is often inferred from very simple parameters (such as lift and drag coefficients and glide angles); these alone may not be sufficient measures of aerodynamic performance because animals flying in real environments will experience perturbations and the need to maneuver around obstacles [1].

Discoveries [2–8] during the last decade of a diversity of feathered dinosaurs and early birds from the Mid-Late Jurassic through the Cretaceous of Liaoning, China have led to considerable speculation about the roles that the feathers played on these extinct animals. Fossil forms are important and informative in biomechanical studies because they may indicate transitional forms within a lineage between ancestral and derived taxa, or they may record natural experiments in form, particularly in side-branches of the tree. Although we cannot observe the behavior of extinct animals, we can measure the aerodynamic forces on dynamically-scaled physical models in a wind tunnel to quantify the broader effects on performance of different postures and morphologies. Since physical laws apply to all taxa, regardless of history, knowing about the physical implications of shape can suggest suitable prior assumptions (for example, plesiomorphies; starting estimates for aerial performance within a clade; other limits based on performance that can be ruled out) that should apply in comparative studies of physically-constrained aerially maneuvering animals of similar shape.

We used physical models [9], based on †*Microraptor gui* (Fig. 1), a cat-sized dromaeosaur with flight feathers on its forelimbs, hindlimbs, and tail. The models enabled us to investigate effects of diverse aerodynamic surfaces in the aft/posterior of a body and of various movements of the appendages. By measuring not just lift and drag, but also side forces and moments in pitch, roll, and yaw, we can assess static aerodynamic stability (tendency to experience righting torques when perturbed) and control effectiveness (moments generated by motions of control surfaces), both of which affect the ability to maneuver while gliding or parachuting through a complex forest habitat [10, 11].

**Figure 1.**
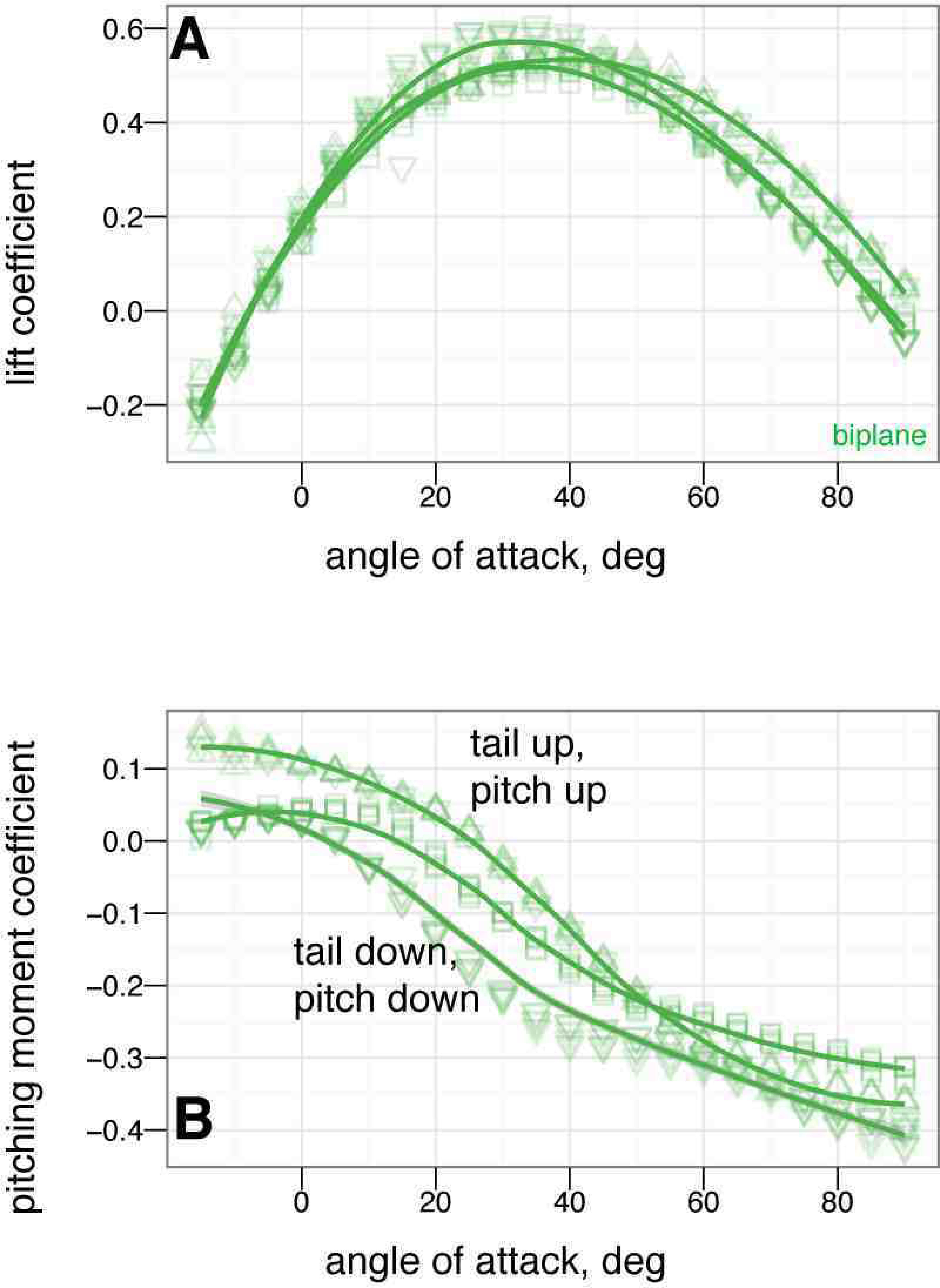
†*Microraptor gui* from [2], a dromaeosaur from the Cretaceous Jiufotang Formation of Liaoning, China; physical models, and sign conventions. A, Holotype specimen IVPP V13352, scale bar 5 cm. Notable features include semilunate carpal bones, a boomerang-shaped furcula, a shield-shaped sternum without a keel, uncinate processes on the ribs, unfused digits, an intermediate angle of the scapulocoracoid, and a long tail of roughly snout-vent length. In addition, there are impressions of feathers on the forelimbs, hindlimbs, and tail. B-J, Physical models of †*M. gui*, scale model wingspan 20 cm, snout-vent-length 8 cm. Reconstruction postures, B-I, used for constructing physical models: B, sprawled, after [2]; C, tent, after [34, 58]; D, legs-down, after [34]; E, biplane, after [32]. F-I additional manipulations: F, asymmetric leg posture with 9090° leg mismatch (*arabesque*); G, example asymmetric leg posture with 45° dihedral on one leg (*dégagé*), H, sprawled without leg or tail feathers; I, tent without leg or tail feathers. J, test setup; K, sign conventions, rotation angles, and definitions for model testing, after [10, 30, 31].

The Jiufotang Formation, in which †*M. gui* was found, has been interpreted as a forest based on pollen data and plant fragments [3, 12]. The inference that †*M. gui* was arboreal solely based on pollen is not terribly strong, given that not everything that lives in a forest lives in the trees and that processes after death (taphonomy) that occur during fossilization also tend to mix the remains of organisms from different habitats together [13]. However, many organisms in forests make use of the trees even if they don’t appear particularly arboreal [14]. In addition, the vertebrate fauna in this formation includes several species of pterosaurs [15,16] as well as numerous feathered theropod dinosaurs and basal (†*Jeholornis* and †*Sapeornis*) and enantornithine birds [3, 17–21]. Gut contents of one extraordinary specimen consisted entirely of arboreal enantornithines [22]. Of the feathered dinosaurs, many are of small size [23, 24] and similar feathered forms with varying degrees of leg- and tail feathers, suggesting that at least some might have been in the trees and performing aerial behaviors; morphological data also suggests arboreality [22, 24]. Therefore, in examining †*M. gui*, it is worthwhile to consider arboreality, aerial hypotheses [1] and the role of aerodynamic forces and torques, rather than constrain thought only to behaviors reliant on ground contact.

### Aerodynamics and the effects of shape and posture: hypotheses

In this paper, we discuss results of a systematic survey of stability and control effectiveness in a four-winged [2, 25, 26] ancestral morphology [8, 27]. Our models are based on one specimen of †*M. gui*, but the four-wing plus feathered tail pattern is now considered to be ancestral to the Avialae [8, 27–29]. In particular, we examine the effect of feathered hind limbs and tail (which we hypothesize had may have functioned as empennage by stabilizing the body or providing control) on stability and control effectiveness, as well as control movements of the feathered fore limb / wings. We hypothesize that shape and posture can affect aerodynamic stability and maneuverability. These effects may be larger and are potentially more relevant to early animal flight performance or flight performance in constrained environments than typical metrics of gliding performance based on lift-to-drag ratios.

To quantify stability, we measured rolling, pitching, and yawing moments on models in different postures and body positions, held at fixed orientations relative to the air flow. The moments were used to examine the slope near fixed points where moments were zero [10,30,31]. For a quasi-static situation, a positive slope means that the resulting moments will tend to increase a directional perturbation, while a negative slope indicates a restoring moment that resists the perturbation. This provides a way to diagnose stability as a three-character trait (positive slope unstable, zero slope marginally stable, negative slope stable). To examine the effect of shape on stability, we measured stability for models with leg and tail feathers versus without leg and tail feathers. We also tested the models in different baseline postures proposed in the literature [2, 32–34].

The control effectiveness of different movements can be measured by deflecting the appendages (forelimbs, hindlimbs, and tail) and measuring changes in the moments. Control effectiveness identifies which appendage movements are effective in creating forces and torques that can be used for maneuvering, and which appendage movements are not effective. When considering the use of wings, such as in flapping fliers like *Calypte anna* or gliders like *Draco*, or other appendages of intermediate function, such as in frogs [10], bristletails [1], stick insects [35], ants [36], or humans [37,38], it becomes clear that a wide range of symmetric and asymmetric movements can be used and that effective movements may vary depending on the flight regime. We hypothesize that symmetric appendage movements, in which left-right pairs of appendages are moved together) will be most effective in pitch, while asymmetric movements, in which left-right differences are created, will create rolling and yawing movements (see also chapter 1 of [39]). Based on intuition from activities like skydiving and windsurfing, the most effective control movements should involve large motions of big surfaces (e.g. long tails or large wings) far from the center of mass. For human skydivers in freefall, several stable and unstable postures are possible. The effectiveness of symmetric movements in controlling pitch and asymmetric movements in generating yaws and rolls was demonstrated in [37, 38]; windsurfers create yaw by protracting or retracting the entire sail relative to the keel center of pressure, using a universal joint roughly comparable to a gleno-humeral joint.

Vertebrate fliers (typically considered to include birds, bats, pterosaurs) have converged on a two-wing geometry with “high” aspect ratio (*s*^2^/*A* > 0.5, wider than a pancake) although larger variation in geometry is seen when considering all vertebrate taxa with aerial behaviors; those other taxa also make wide use of various body parts to accomplish maneuvers [1, 40]. In particular, the multiple feathered surfaces of †*Microraptor* might be expected to have large impacts on maneuvering [41–44]. Multiple control surfaces may have important functional consequences. For example, in engineering practice, rolls up to large angles in submarines can be caused by interactions between the sail (upstream appendage) and rudder (downstream appendage). In the submarine case, dihedral planes are sometimes added to stabilize the ship; we hypothesize here that leg feathers may have such a stabilizing role. To consider a living example, interactions between median or paired fins can enhance maneuvering in fish [41,42,45,46]. A four-(or more) fin planform is widely seen in aquatic creatures, and also occurs in some “gliders” like frogs [10] and four-winged flying fish [47]. Multiple tandem aerodynamic surfaces can also result in delayed onset of stall.

Finally, we wish to test if the function of appendages varies with the aerodynamic environment. In other studies, fluid environmental characteristics such as Reynolds number (Re, a nondimensional measure of the relative importance of inertial to viscous effects) can result in shifts in the function of an appendage [48, 49]. In this study, vertebrate fliers are large, fast, and at high angle of attack, turbulent. Rather than Re [50], more important parameters for flight may be the angle of attack or glide angle. Angle of attack (relative to oncoming airflow) and glide angle (relative to horizontal) are not the same, but many animals with high glide angle aerial behaviors are also at high angle of attack [1, 36–38]. Speed, angle of attack, and glide angles are kinematic variables that may be expected to change as aerial behaviors evolve. Directed aerial descent performance at high glide angles and angles of attack is widley distributed, even among taxa without obvious aerial features [1, 35, 36], and is possible even in vertebrates [37, 38, 40]. During a transition between high glide angle directed aerial descent and lower angle behaviors, the function of appendages in creating aerial forces and moments may shift, or completely reverse behavior. In engineering practice, this phenomenon is termed “reversal”. As an example, ship rudders at low speed act can act opposite to their normal behavior. Helmsmen unaware of such phenomena have caused collisions. High angle of attack aerodynamics may be different from low angle of attack in important ways for organisms in the process of evolving flight, especially when such shifts in stability and control effectiveness are considered. We hypothesize that shifts in stability and control effectiveness are linked to the angle of attack while the effect of Re will be small in comparison (although it is good practice to check for scale effects in any model test.)

### Review of previous model tests in dinosaurs

Tests using dynamically similar models of animal shapes have long been done; Reynolds’ original work included ducks [9]. Dinosaur flight mechanics have been previously studied using both computational and experimental approaches. Generally, fluid mechanics benefits from use of both approaches.

Heptonstall [51] examined †*Archaeopteryx*, and later Gatesy and Dial [52] examined †*Archaeopteryx* tails using computational approaches, both without benchmarking against experiment. Longrich [53] later recognized the presence of leg feathers in †*Archaeopteryx* and provided the first estimates of dinosaur maneuvering capabilities via computations based on [10, 54]. Chatterjee and Templin [55] used computer simulations for assumed aerodynamic coefficients to identify phugoid mode gliding in †*Archaeopteryx*; these were later extended to a particular biplane configuration of †*Microraptor* [32]. These are computational studies, using coefficients and assumptions drawn from fixed wing aircraft at low angle of attack. In particular, early vertebrate fliers may not be using low angles of attack [1, 39, 56, 57], and long glides or high *L/D* may not be driving evolution of flight [1, 35, 36, 39].

Model tests have been used in more recent dinosaur studies. Xu, Jenkins, Breuer, et al. used full-scale wind tunnel models constructed by professional preparators to examine flight characteristics of †*Microraptor* (Provided in a TV documentary in [34]; data not yet published; leg positions described in [58]). The results of that program focused on lift and drag and only briefly addressed stability. The methods here are most similar to that effort, and to recently published results from [57] as well as computational results from [59].

Alexander et al. [33] also used full-scale flying models constructed from styrofoam gliders, to test the biplane hypothesis of [32]. While we agree that models can provide useful aerodynamic information, we note that additional nose ballast was needed to allow stable flight in the chosen wing configuration. We, and others [60], are unsure what is the anatomical basis for creating a stiff lower-wing with feathers cantilevered from the tarsometatarsus. As there are results both in support of [33, 57] and against [60] this posture, we tested it.

### Limitations of the model testing approach

In reconstructing the biomechanics of extinct animals, it is important to restate several caveats. We consider the largest source of uncertainty in these types of studies to be the use of one (or a few) specimens; this is a common uncertainty in many fossil aerodynamic studies, especially those based on †*M. gui* [32–34] and others. Simply put, the model is never known to the same precision as a production aircraft or airfoil section and, while it is important to learn what we can from fossils, we should be careful not to over-reach with partial data from small numbers of specimens.

The next largest source of uncertainty is the reconstruction shape and posture; our response here is to test many proposed reconstructions and examine the functional consequences. The remaining sources of uncertainty include variation in placement of the model on the supporting sting and in positioning and construction of individual models; to show the bounds of these, we have plotted all replicates and included all runs, including those with small misalignments.

Another limitation of model tests that must be acknowledged is that the live animal may have used closed-loop neuromuscular control or engaged in movements that would result in more dynamic behaviors. The results here may still be applicable to this case for several reasons. In many animal movements, including in flight, passive stability is exploited where available. Many “unsteady” flight movements can still be modeled as quasi-static (as in the simulations of [32, 57, 59], or in our other work). In addition, closed loop control can only make use of a control mechanism when it actually has some amount of control effectiveness (those with no control effectiveness can be ruled out as means for effecting maneuvers). For the advance ratio approaching forward flight (*J* = *u/*2*ϕnR* ∼ 4), we have also constructed flapping models (in preparation as a separate paper). For the most dynamic flapping situations, we advocate live animal studies [36, 39], as well as freely flying models (see also [61–64]).

## Results

During the fall of 2010, we collected a dataset of 12,810 measurements for 180 combinations of postures and positions, with at least five replicates for each. The raw data require approximately 5.3 GB of storage. The work was accomplished during approximately 350 hours of wind tunnel time by a team of ten undergraduates led by one graduate student. Reduced data has been deposited on the public repository Bitbucket (https://bitbucket.org/devangel77b/microraptor-data), and can be downloaded, along with all R code, as a zipped archive at: https://bitbucket.org/devangel77b/microraptor-data/downloads/microraptor-data.tar.gz

For the plots given here, color represents the base posture: red for sprawled, blue for tent, green for biplane, and purple for down. All sign conventions are as in [10, 30, 31] and as shown in Fig. 1. Symbols, where used, represent variations in position from the base posture, such as movement of legs, wings, or tail. All units are SI unless otherwise noted. To standardize comparisons for speed and a baseline posture (adopted from [2]), raw forces and torques were nondimensionalized into aerodynamic coefficients (e.g. lift and drag force coefficients of the form *F*/0.5*ρU*^2^*A*, and moment coefficients of the form *M*/0.5*ρU*^2^*Aλ*) as outlined in Methods.

### Baseline longitudinal plane aerodynamic data and effects of posture and the presence/absence of leg and tail feathers

Fig. 2 gives the nondimensional coefficients of lift, drag, and pitching moment for †*M. gui* with full feathers.

**Figure 2.**
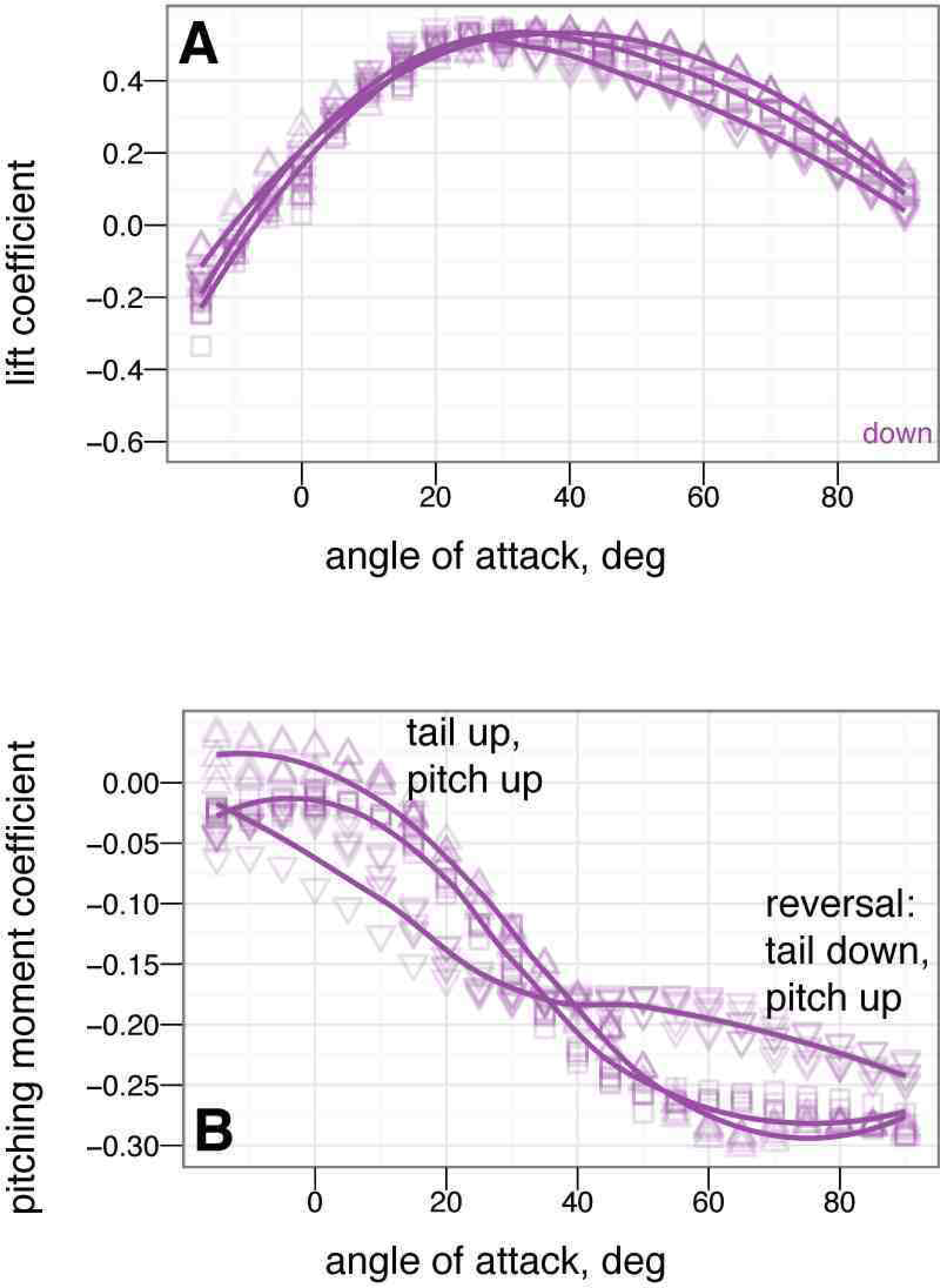
Nondimensional coefficients for all baseline postures. Red is sprawled, blue is tent, green is biplane, purple is down. *α* from –15° to 90° in 5° increments, with five or more replicates per treatment. A, Lift coefficient. B, Drag coefficient. C, Lift drag polars. D, Pitching moment coefficient. Stable angles of attack, which cross *C_m_* = 0 with negative slope, for tent (blue) and biplane (green) postures identified with yellow arrows.

Scaling with the coefficients, the full scale forces for †*M. gui* at 12 ms^−1^ are plotted in Fig. 3.

**Figure 3.**
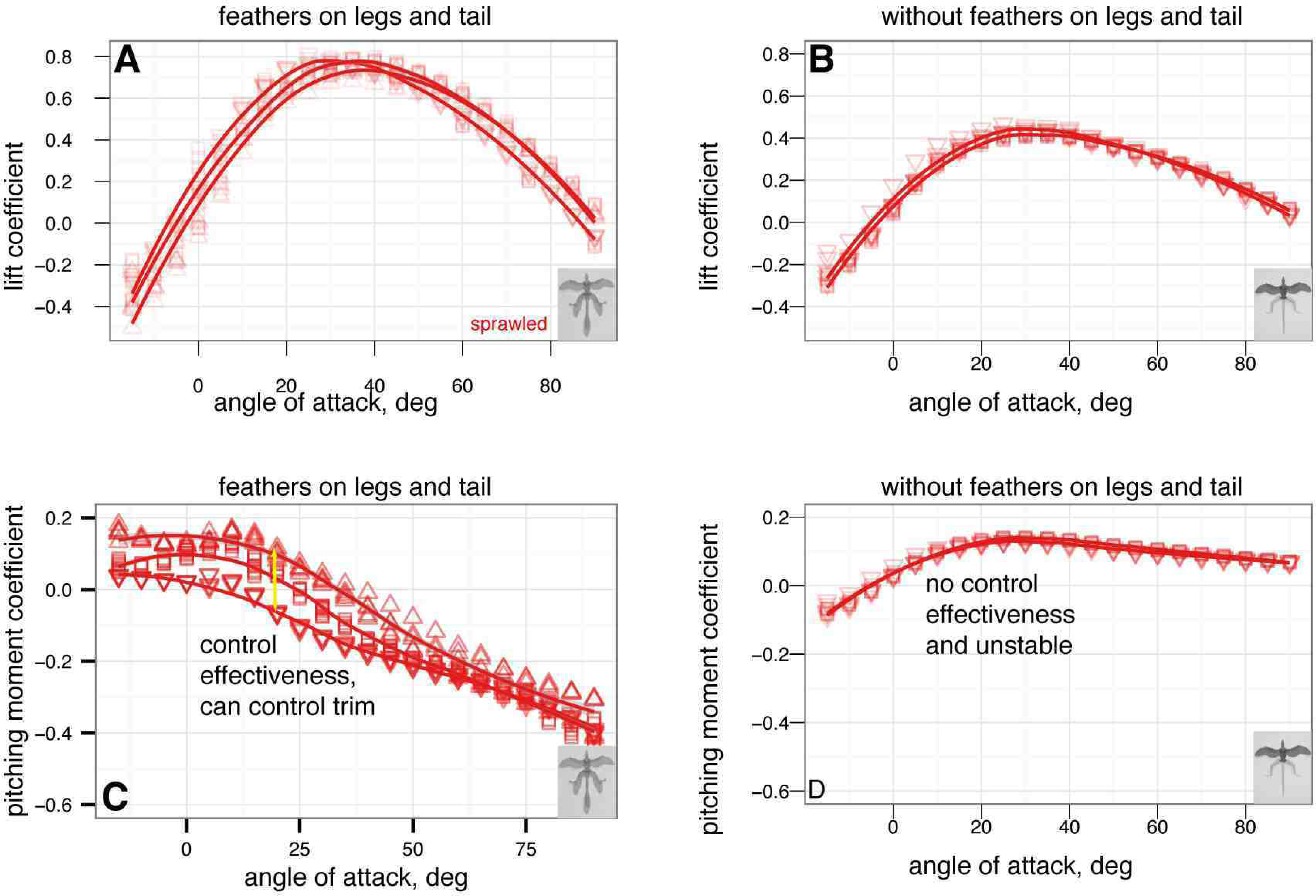
Full scale forces and moments for †*M. gui* at 12 m s^−1^. Red is sprawled, blue is tent, green is biplane, purple is down. *α* from –15° to 90° in 5° increments, with five or more replicates per treatment. Gray band indicates weight range of †*M. gui*. A, Full scale lift at 12 m s^−1^, all models. B, Full scale drag at 12 m s^−1^, all models. C, Lift-drag polars. D, Full scale pitching moment at 12 m s^−1^ versus angle of attack, all models. Stable angles of attack for tent (blue) and biplane (green) indicated.

For comparison with previous work [65], various other gliding performance metrics are compared in Figs. S1 and S2 (available online).

A Reynolds number sweep from 30,000–70,000 (Fig. 4, Table 1) was also conducted to check for scale effects.

**Figure 4.**
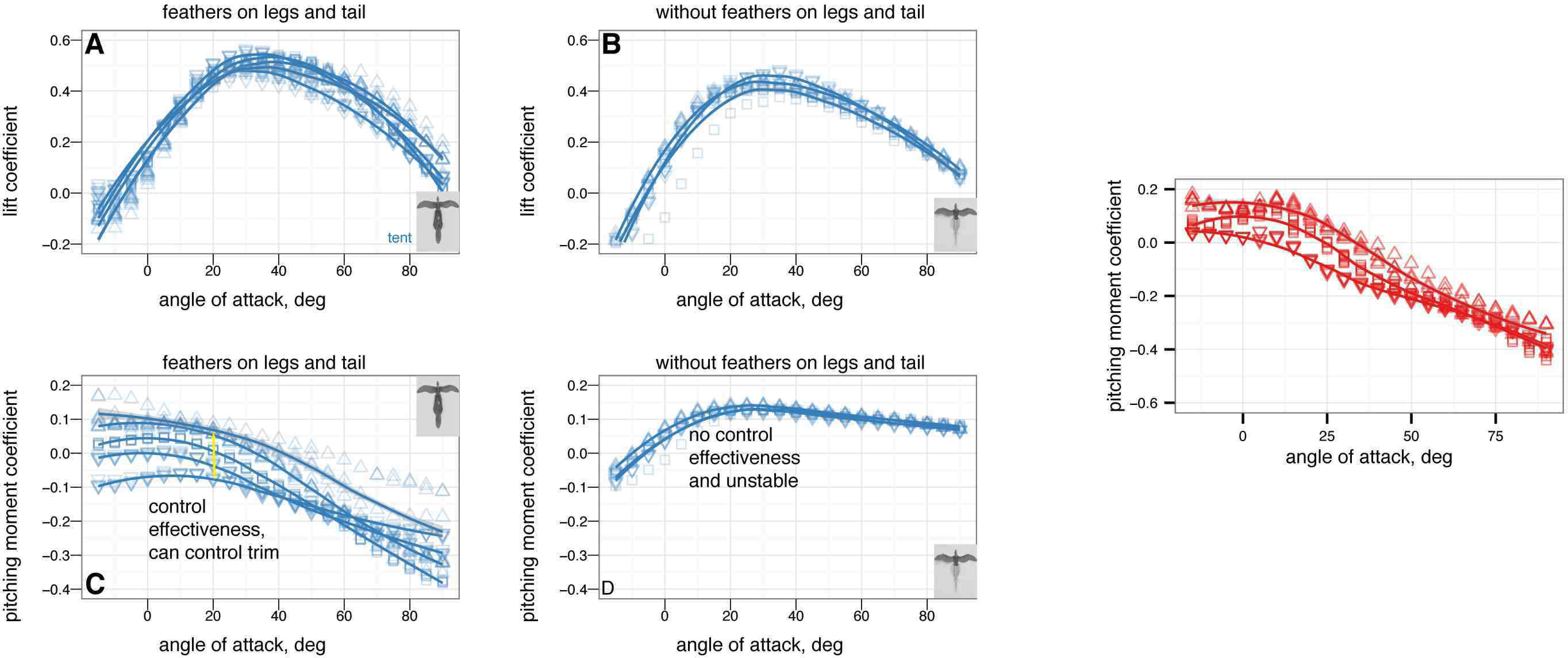
Reynolds number sweeps for A, lift, B, drag, and C, pitch coefficients. There are not large changes in aerodynamic coefficients over the ranges shown here. This is similar to what is seen in benchmarking tests with *Draco* lizard and Anna’s Hummingbird (*Calypte anna*) models. The coefficients are roughly constant in the range of †*Archaeopteryx*. Moment coefficients are constant over the range shown.

**Table 1.**
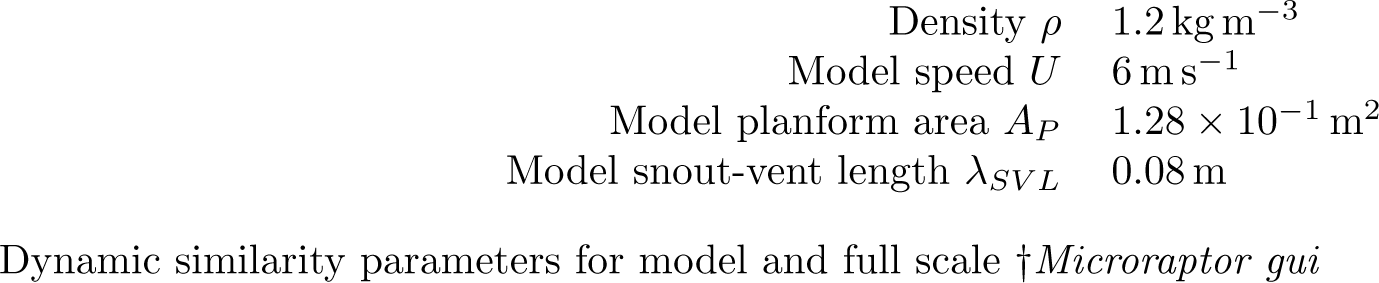
Dynamic similarity parameters

### Effect of leg and tail feathers

The effects on longitudinal plane coefficients of the presence or absence of leg and tail feathers are shown in Figs. 5 and 6.

**Figure 5.**
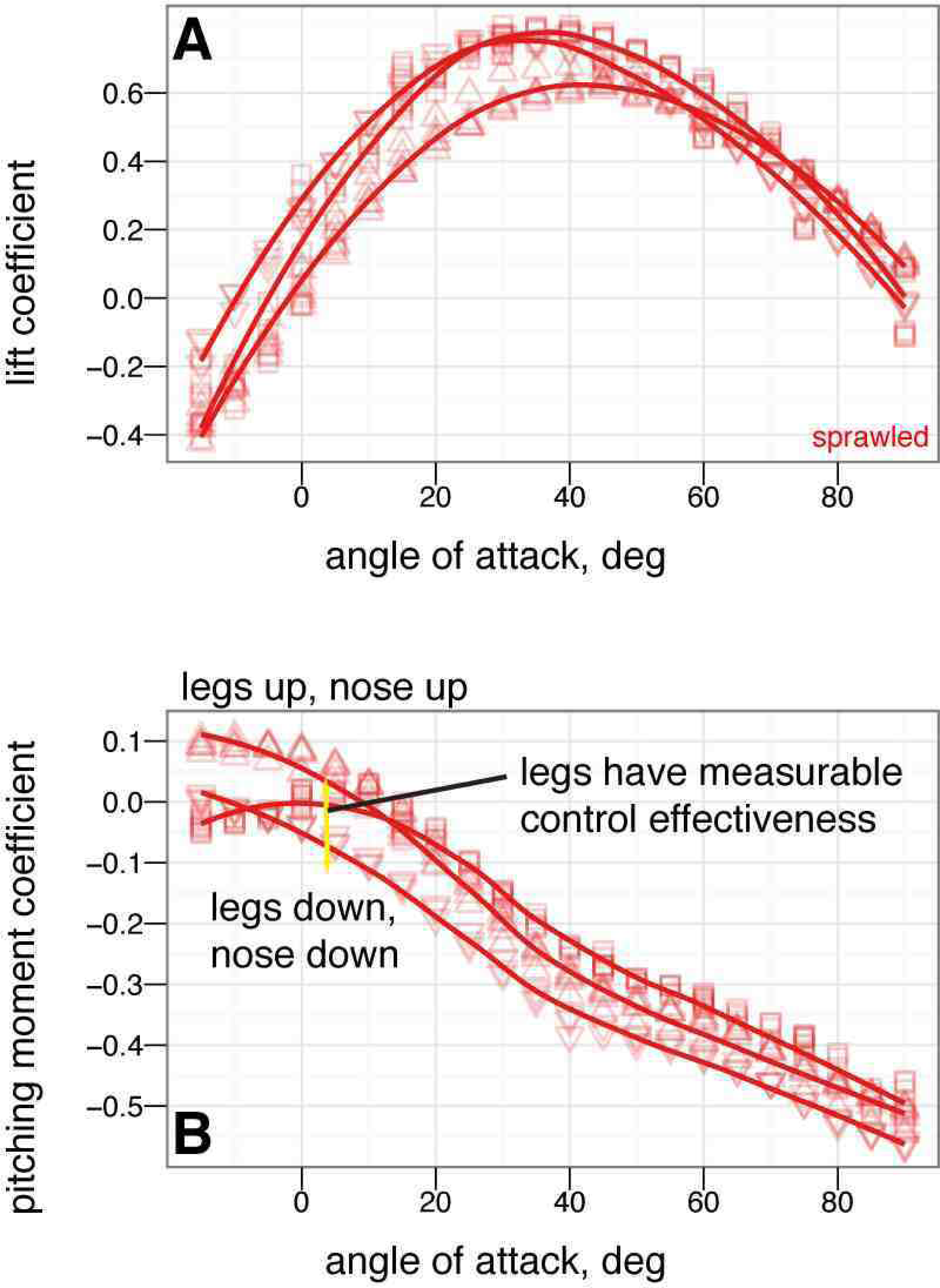
Presence or absence of leg and tail feathers can substantially alter longitudinal plane aerodynamics. Sprawled and tent postures with and without feathers, all coefficients shown versus angle of attack, solid squares with leg and tail feathers, open squares without leg or tail feathers. A, Lift coefficient. Stall occurs at higher angle of attack when leg feathers are present. B, Drag coefficient. Leg feathers increase drag at high angle of attack, improving parachuting performance. C, Lift coefficient versus drag coefficient. D, Lift to drag ratio. Lift to drag ratio is improved slightly without the additional drag and less-efficient lift generation of hind wings. E, Pitching moment coefficient. Without leg feathers, stability is not achieved in either posture. F, Pitching stability coefficient.

**Figure 6.**
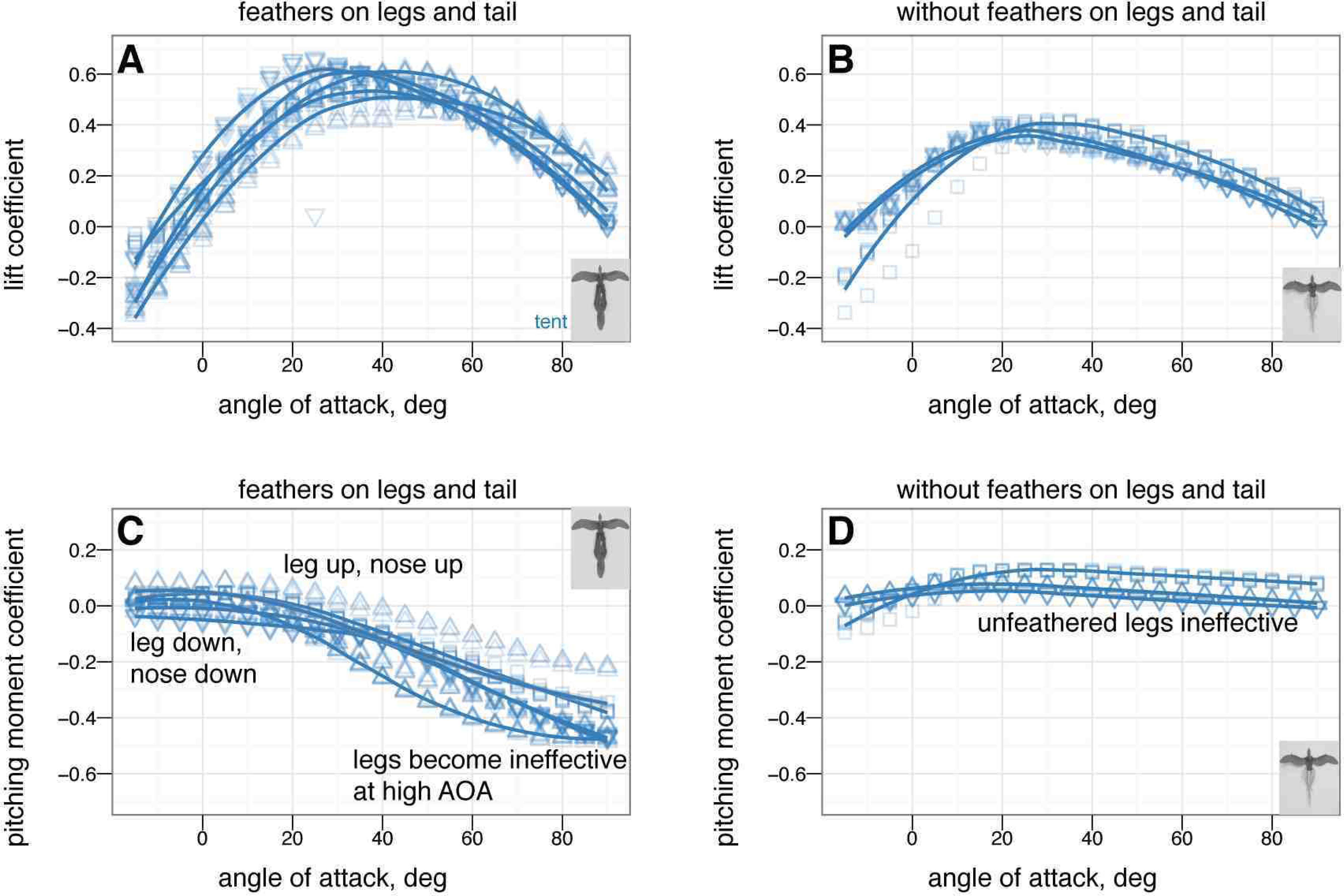
Presence or absence of leg and tail feathers has effects on [65] metrics, although the usefulness of [65] is questionable (see Fig. S2). Feathers present (black outline) or absent (grey outline) A, Maximum lift to drag ratio, by sprawled and tent postures with and without feathers. The maximum lift to drag ratio for tent without leg or tail feathers is significantly higher than for other postures (ANOVA, *P <* 0.003), however, this improvement is never achieved because the tent posture is never stable without leg feathers. B, Minimum glide speed, by sprawled and tent postures with and without feathers. There are no differences in minimum glide speed between postures (ANOVA, *P >* 0.08). C, Parachuting drag, by sprawled and tent postures with and without feathers. There are significant differences in parachuting drag between postures (ANOVA, *P <* 0.04), however, the straight-down parachuting position is not stable.

### Yaw stability and the effects of shape and angle of attack

Fig. 7 shows how yaw stability varies between postures. To examine the effect of aerodynamic environment (*vis-a-vis* glide angle, or angle of attack as a loose proxy for glide angle), Fig. 8 shows how yaw stability changes as angle of attack increases from 0° to 60° to 90°, or how yaw stability would change in going from falling from a tree at high angle of attack, after a launch or jump, to gliding at a low angle of attack. The presence or absence of leg and tail feathers (Fig. 9) also alters yaw stability [66].

**Figure 7.**
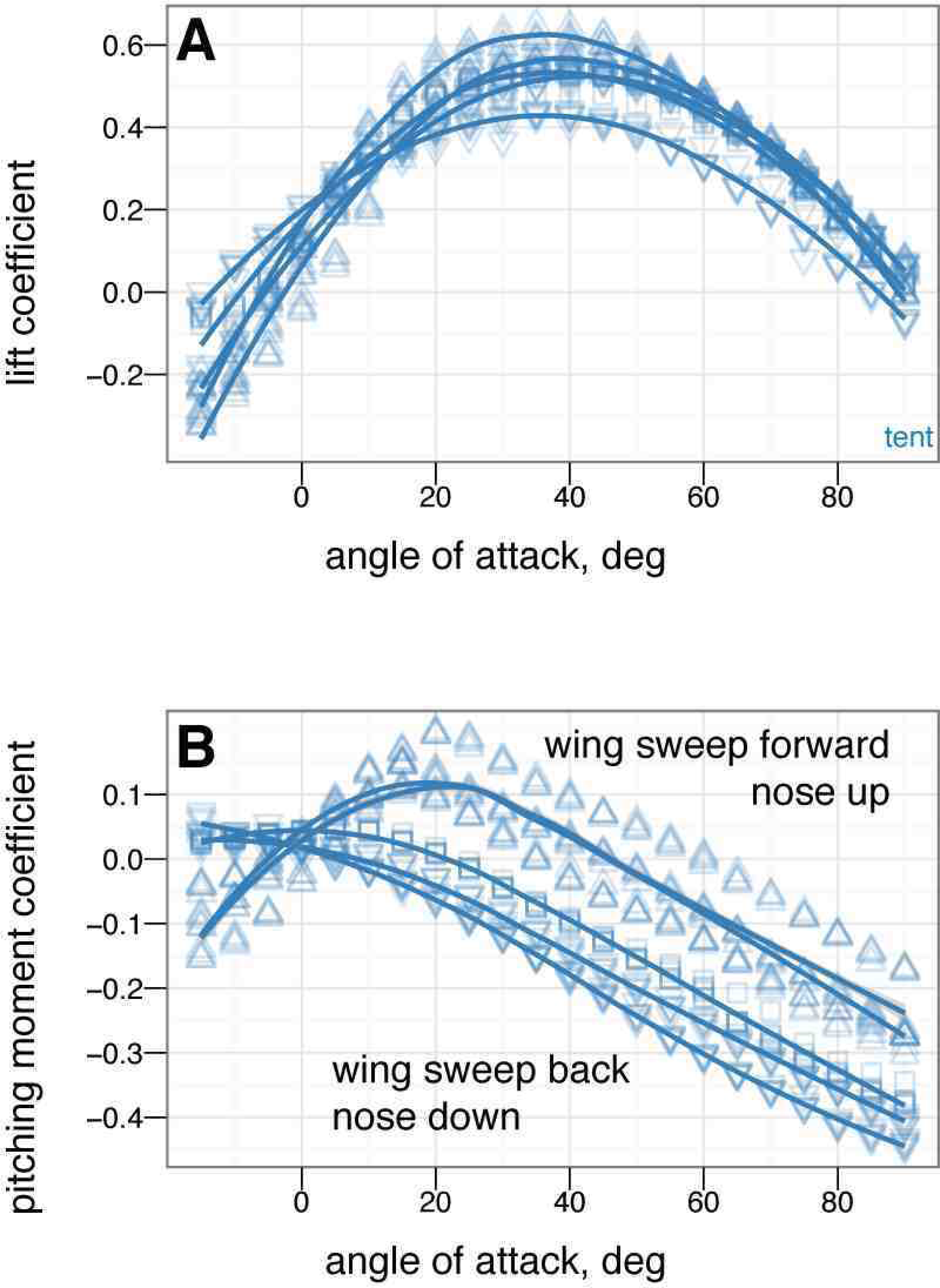
At 0° angle of attack, there are clear differences in yaw stability between postures. In particular, with legs down, the legs strongly act as weathervanes to stabilize the body in yaw (purple line, high slopes near 0°). Color represents the base posture: red for sprawled, blue for tent, green for biplane, and purple for down.

**Figure 8.**
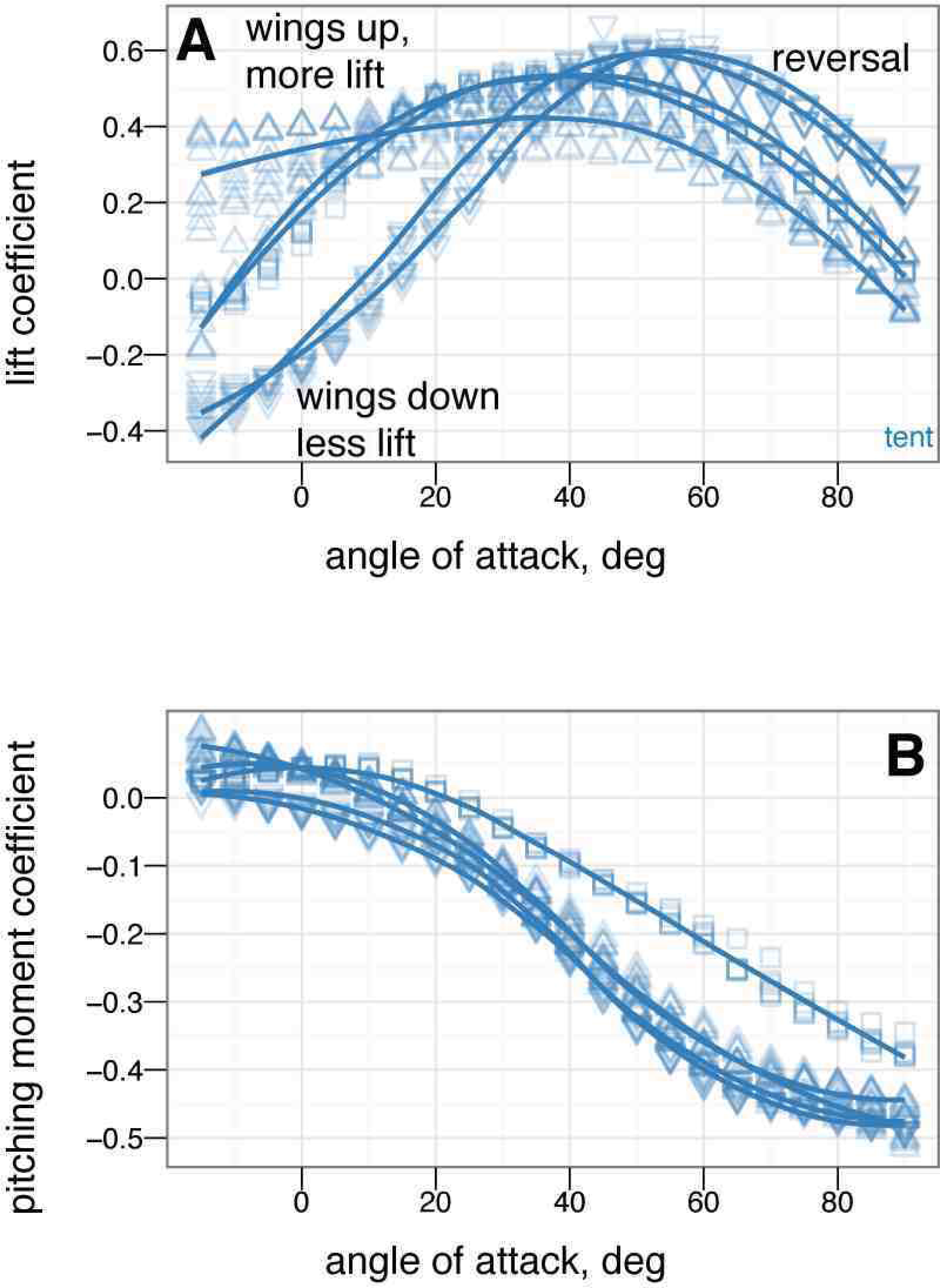
There are also clear differences in yaw stability at different angles of attack. A, At 0°, some postures are more stable in yaw than others. B, At 60°, postures that were stable at 0° may go unstable, such as tent posture. C, At 90°, all postures are marginally stable due to symmetry (lines flat, yawing does not alter position relative to flow). Color represents the base posture: red for sprawled, blue for tent. Organisms may have navigated this transition from 90° to 0°.

**Figure 9.**
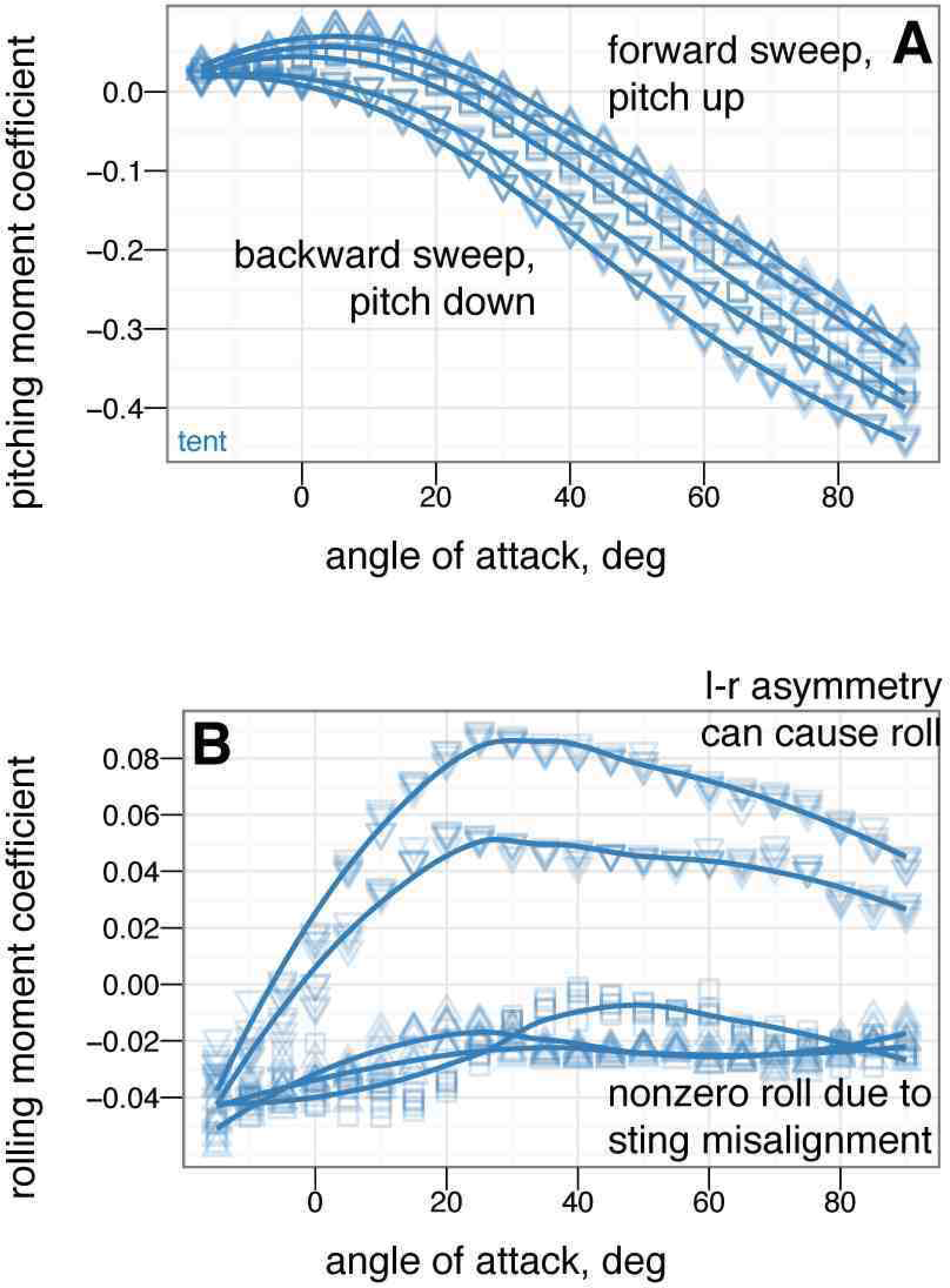
The differences in yaw stability at different angles of attack also depend on the presence or absence of leg feathers. A, At 0°, some feathered-leg postures are more stable in yaw than others. B, At 60°, postures that were stable at 0° may go unstable, such as tent posture with leg feathers. C, At 90°, all postures are marginally stable due to symmetry. Color represents the base posture: red for sprawled, blue for tent, green for biplane, and purple for down.

### Control effectiveness of tail, symmetric wing and leg movements

The control effectiveness for symmetric movements of several appendages is given in Figs. 10 through 17. Figs. 10–13 give the control effectiveness of dorsoventral tail flexion (bending tail up or down 15°) for biplane, down, sprawled, and tent posture. Figs. 14 and 15 give the control effectiveness of symmetric leg movement, in which both legs are deflected, as a pair, in pitch up or down 15°. Fig. 16 gives the control effectiveness for symmetric wing fore-aft sweep (protraction and retraction), in which the fore limb / wings are swept as a pair forwards or backwards up to 45°. Fig. 17 gives the control effectiveness for symmetric wing pronation/supination, in which the wings, as a pair, are pitched up or down up to 30°.

**Figure 10.**
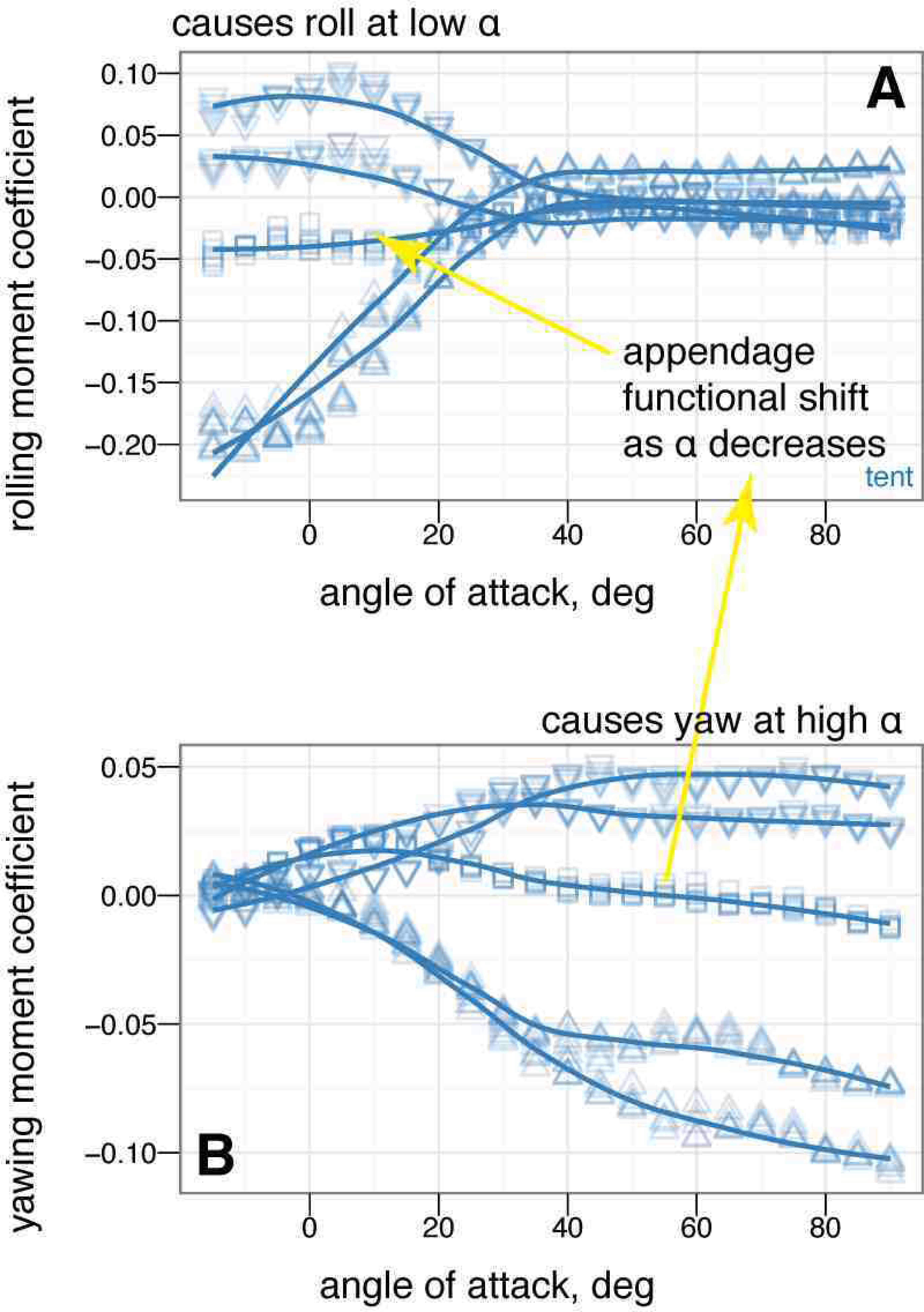
Tail control effectiveness for biplane posture for tail angles of –15° (down triangle), 0° (square), and +15° (up triangle). At low angle of attack, tail up produces a nose up moment relative to zero tail angle, while tail down produces a nose down moment relative to zero tail angle. Tail movement is effective in trimming, by moving the point where the curve crosses *C_m_* = 0. The small effect on lift suggests the tail is primarily effective because of moments generated by its long length.

**Figure 11.**
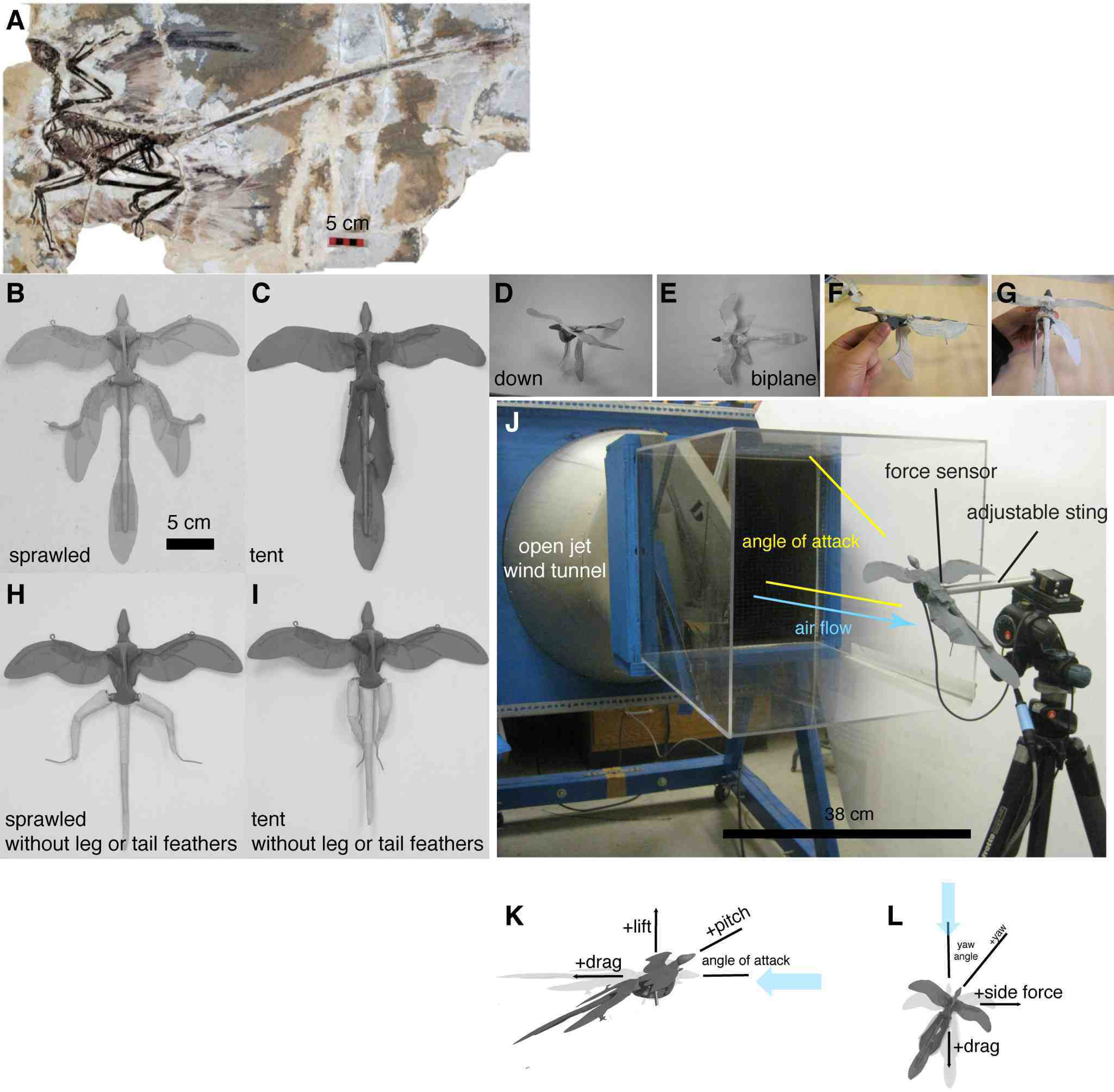
Tail control effectiveness for down posture for tail angles of –15° (down triangle), 0° (square), and +15° (up triangle). At low angle of attack, tail up produces a nose up moment relative to zero tail angle, while tail down produces a nose down moment relative to zero tail angle. Trimming to pitch stability with the tail is only possible with large 15° tail movement. At high angle of attack, the tail experiences reversal in which tail down produces nose up moments/tail up produces nose down moments.

**Figure 12.**
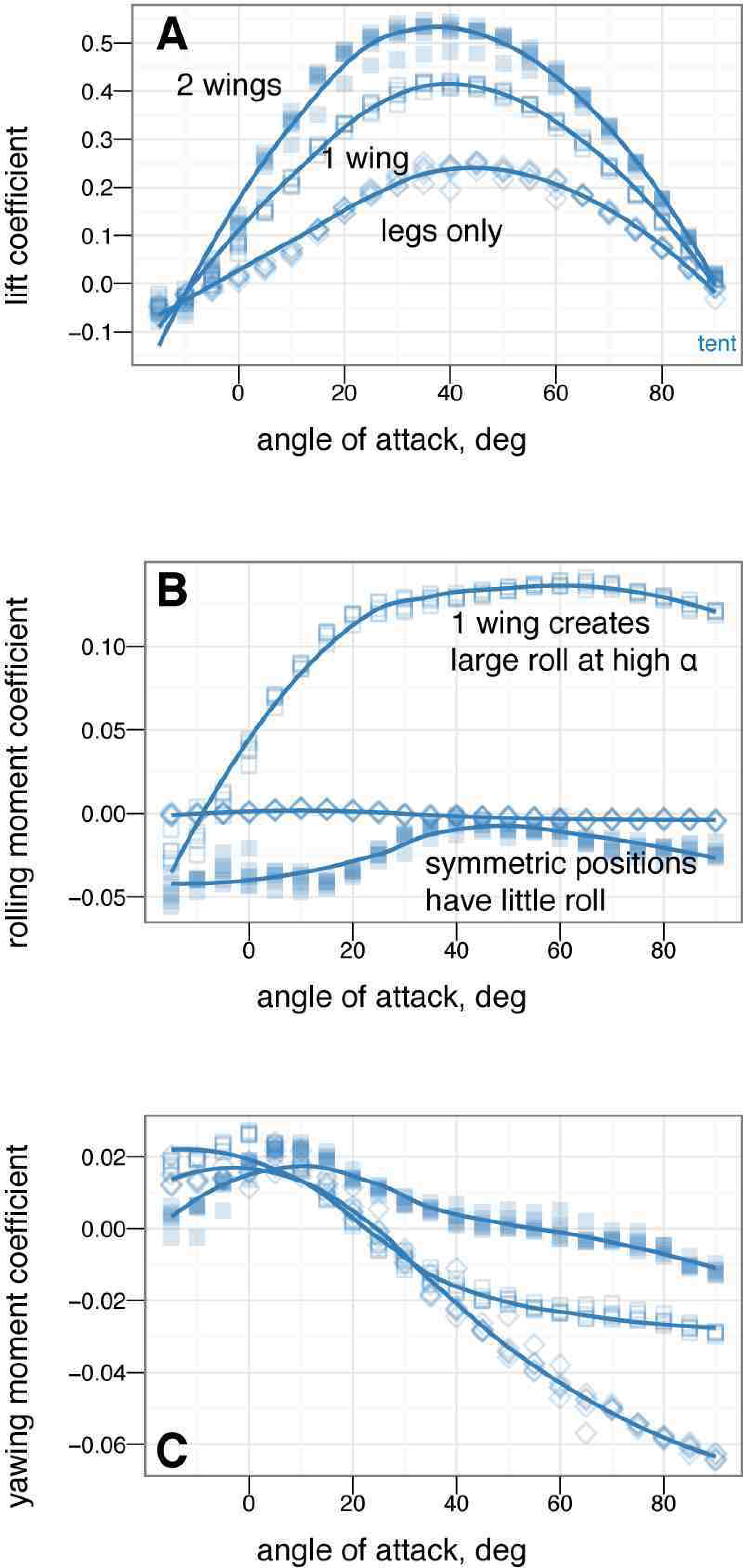
Tail control effectiveness for sprawled posture for tail angles of –15° (down triangle), 0° (square), and + 15° (up triangle), with leg and tail feathers, A & C, and without, B & D. At low angle of attack, tail up produces a nose up moment relative to zero tail angle, while tail down produces a nose down moment relative to zero tail angle, C. Trimming with the tail is able to alter stability. Reversal is not seen at high angle of attack. Without leg feathers, D, the tail is ineffective at producing lift or pitching moment.

**Figure 13.**
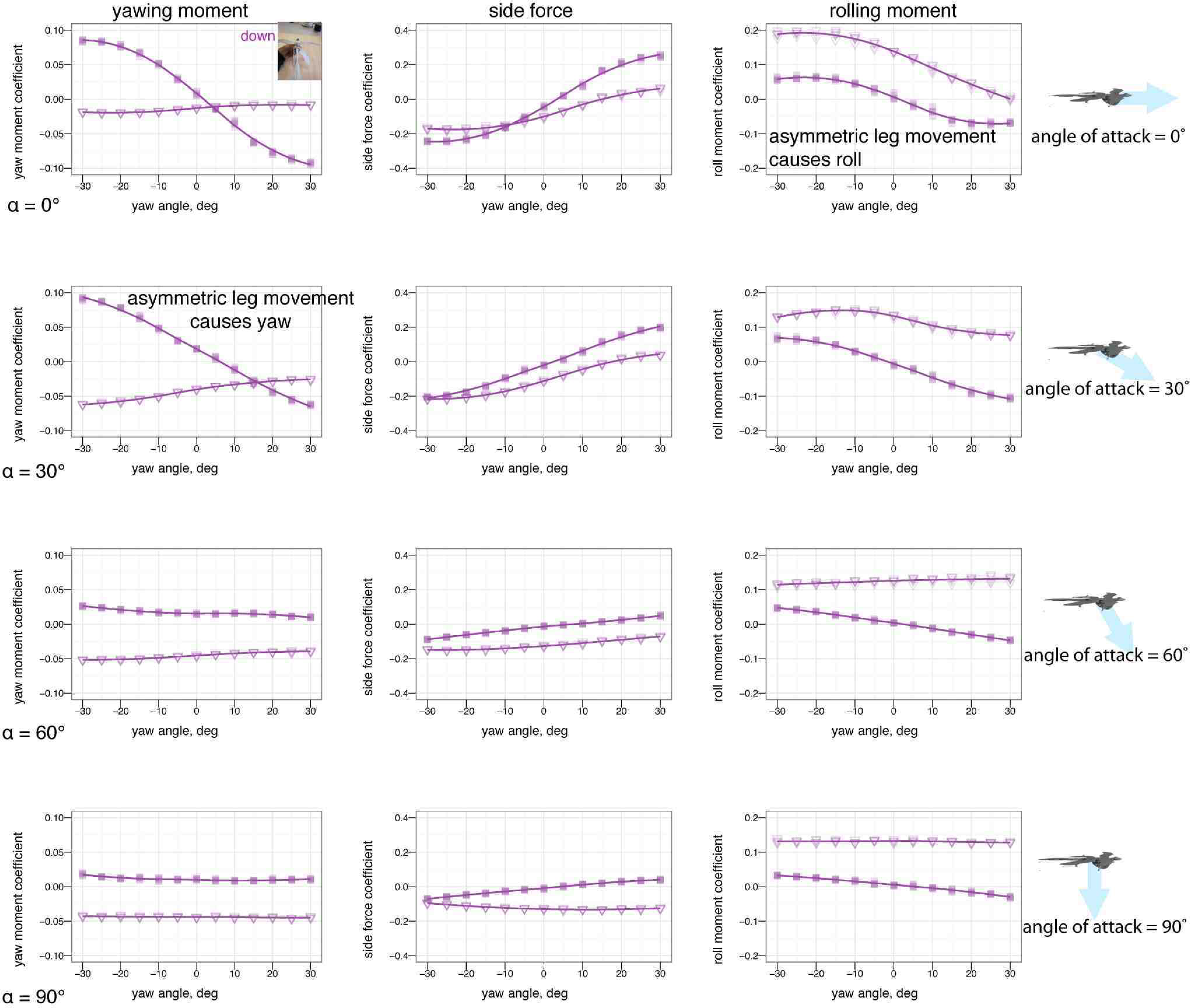
Tail control effectiveness for tent posture for tail angles of –30° (large down triangle),–15° (down triangle), 0° (square), +15° (up triangle), and + 30° (large up triangle), with, A & C, and without, B & D, leg or tail feathers. At low angle of attack, tail up produces a nose up moment relative to zero tail angle, while tail down produces a nose down moment relative to zero tail angle, C. Trimming with the tail is able to alter stability. Some reversal occurs at high angle of attack. Without leg feathers, the tail is ineffective at producing lift or pitching moment, B & D.

**Figure 14.**
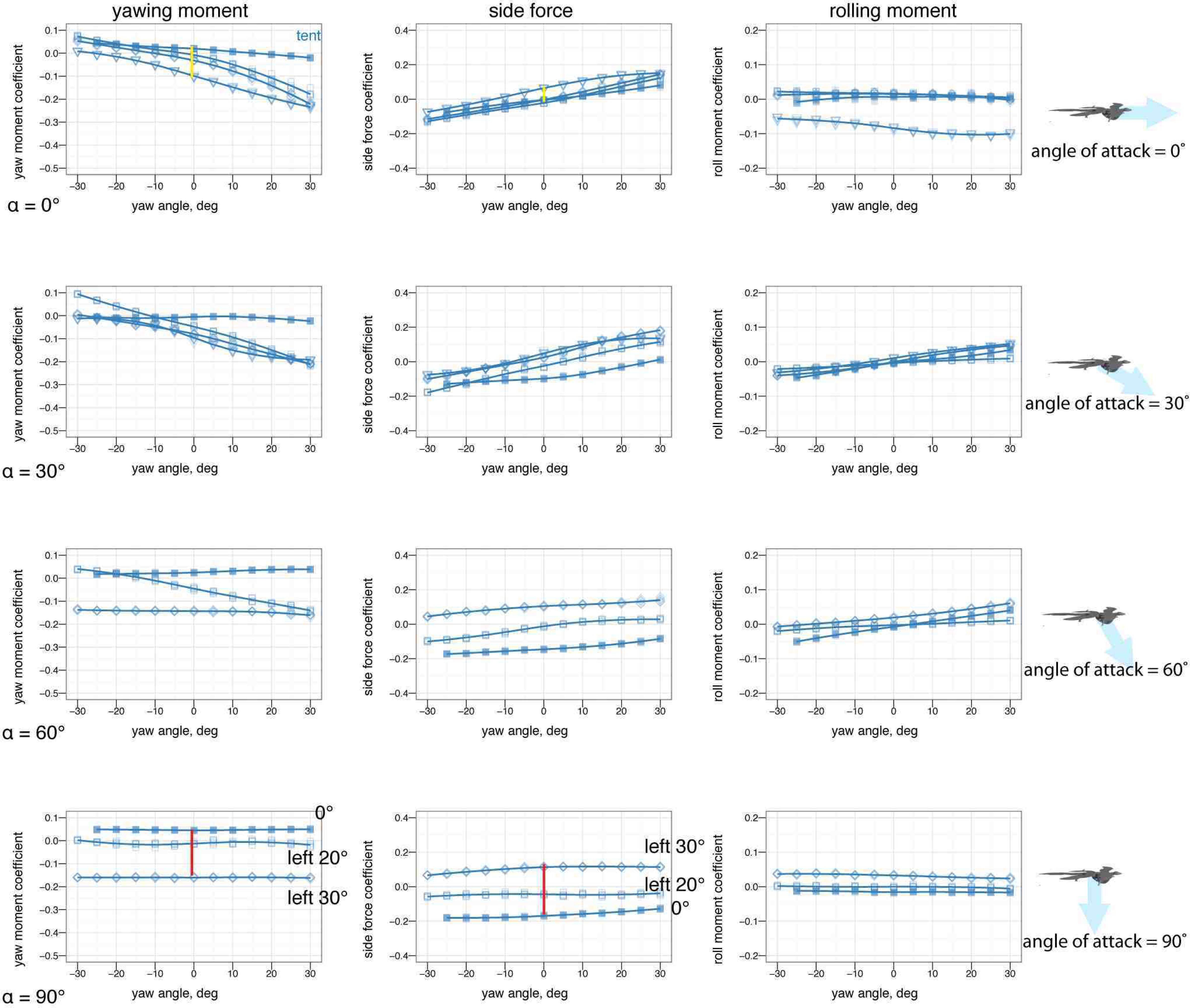
Leg control effectiveness for sprawled posture for leg angles of –15° (down triangle), 0° (square), and + 15° (up triangle). At low angle of attack, legs up produces a nose up moment relative to zero leg angle, while legs down produces a nose down moment relative to zero leg angle. Leg movement is slightly less effective at high angle of attack, and slightly less effective than tail movement.

**Figure 15.**
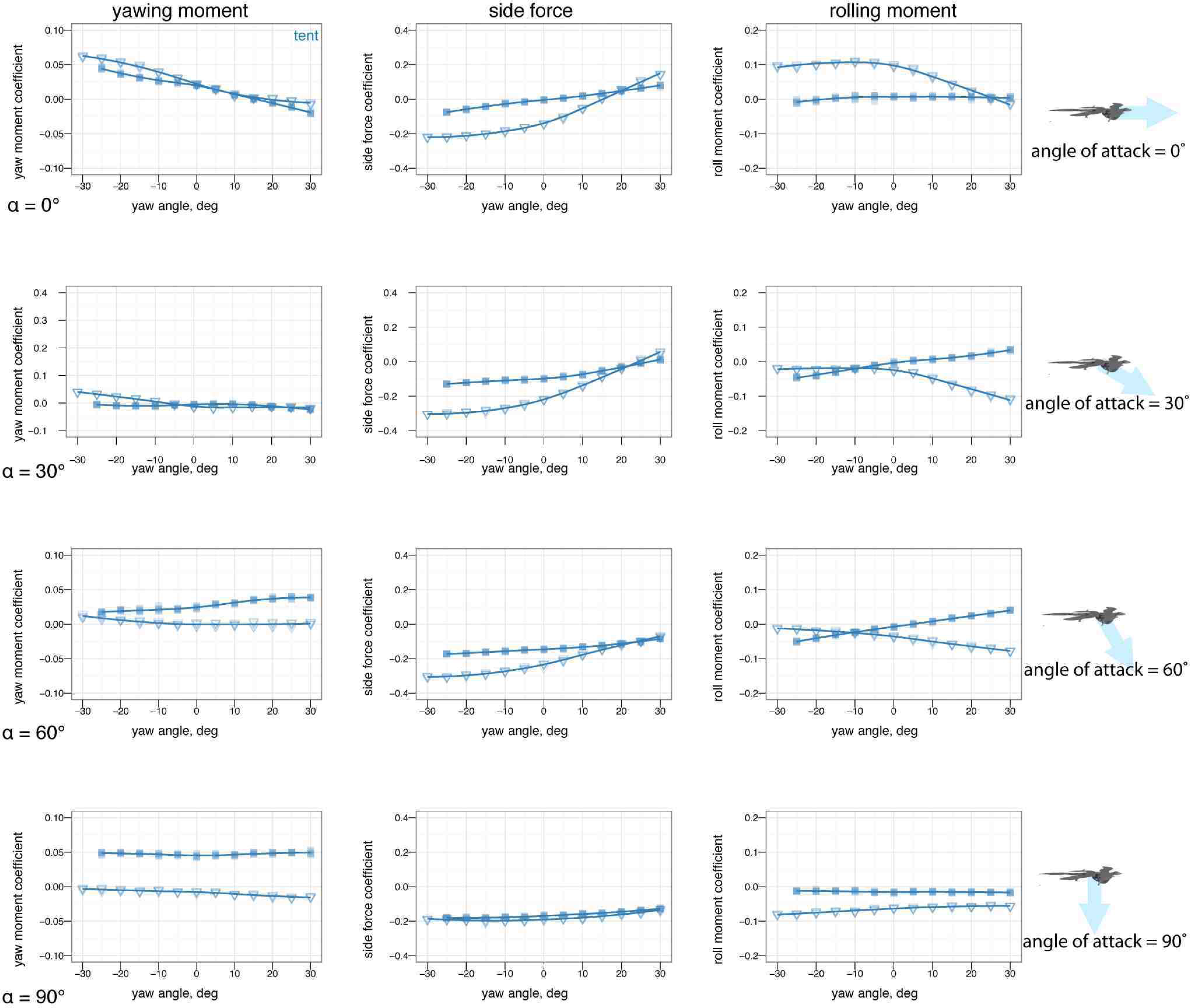
Leg control effectiveness for tent posture for leg angles of –30° (large down triangle),–15° (down triangle), 0° (square), +15° (up triangle), and + 30° (large up triangle) with leg and tail feathers, A & C, and without, B & D. At low angle of attack, leg up produces a nose up moment relative to zero leg angle, while leg down produces a nose down moment relative to zero leg angle, C. Without leg feathers, the legs still have smaller effects, D. At high angles of attack, leg pitch effects become noisy and difficult to identify.

**Figure 16.**
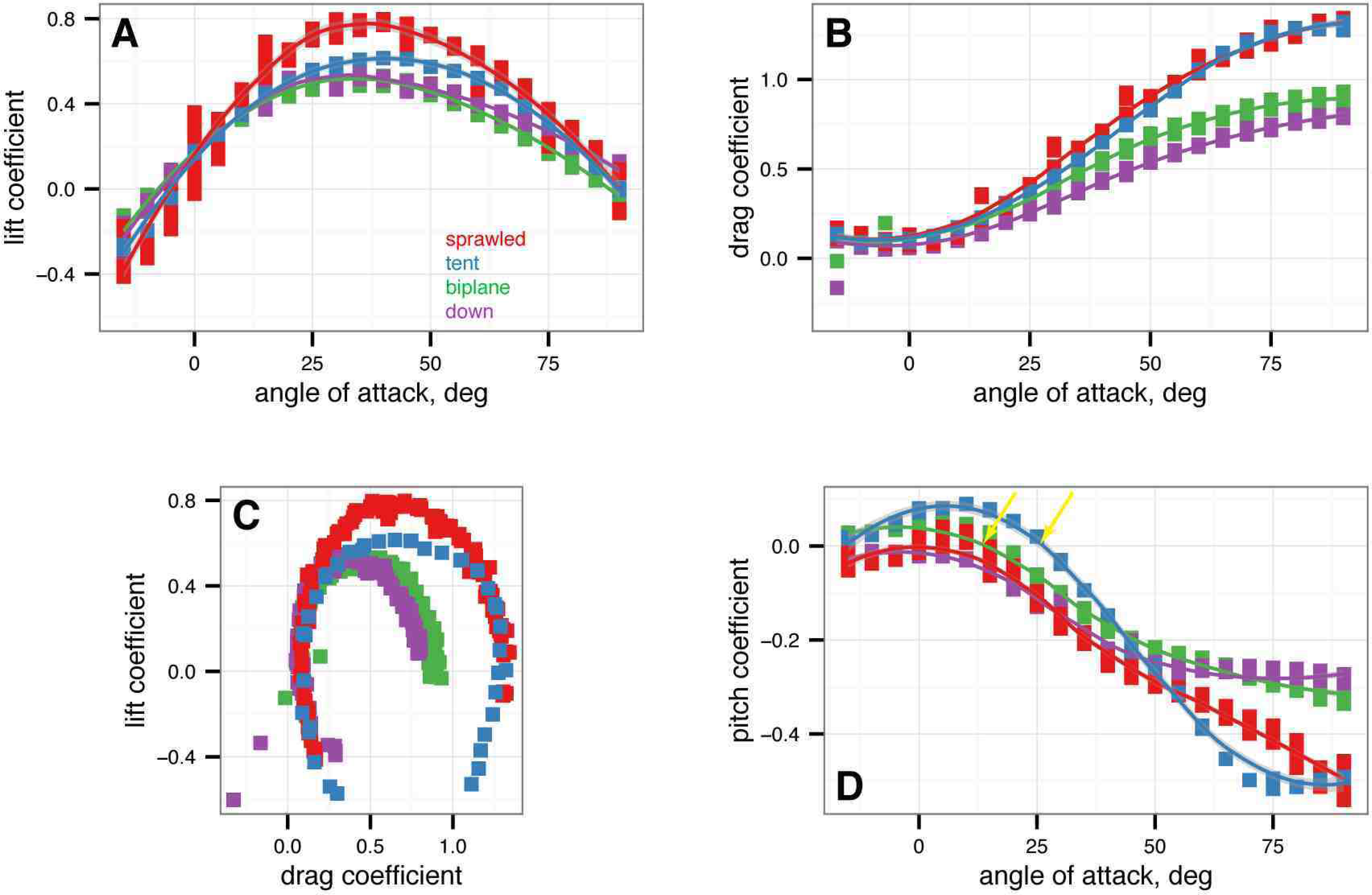
Symmetric wing sweep control effectiveness for tent posture for wing sweep angles of –45° (large down triangle), –22.5° (down triangle), 0°(square), +22.5° (up triangle) and + 45° (large up triangle). Wing sweep is very effective at generating pitching moments. Forward sweep generates nose up moments, while backwards sweep generates nose down moments. This is like steering a wind surfing rig and is similar to what is seen in Anna’s Hummingbird (*Calypte anna*) dive models (Evangelista, in preparation). This mode of control exhibits reversal at negative angle of attack and thus may be difficult to use around 0° angle of attack.

**Figure 17.**
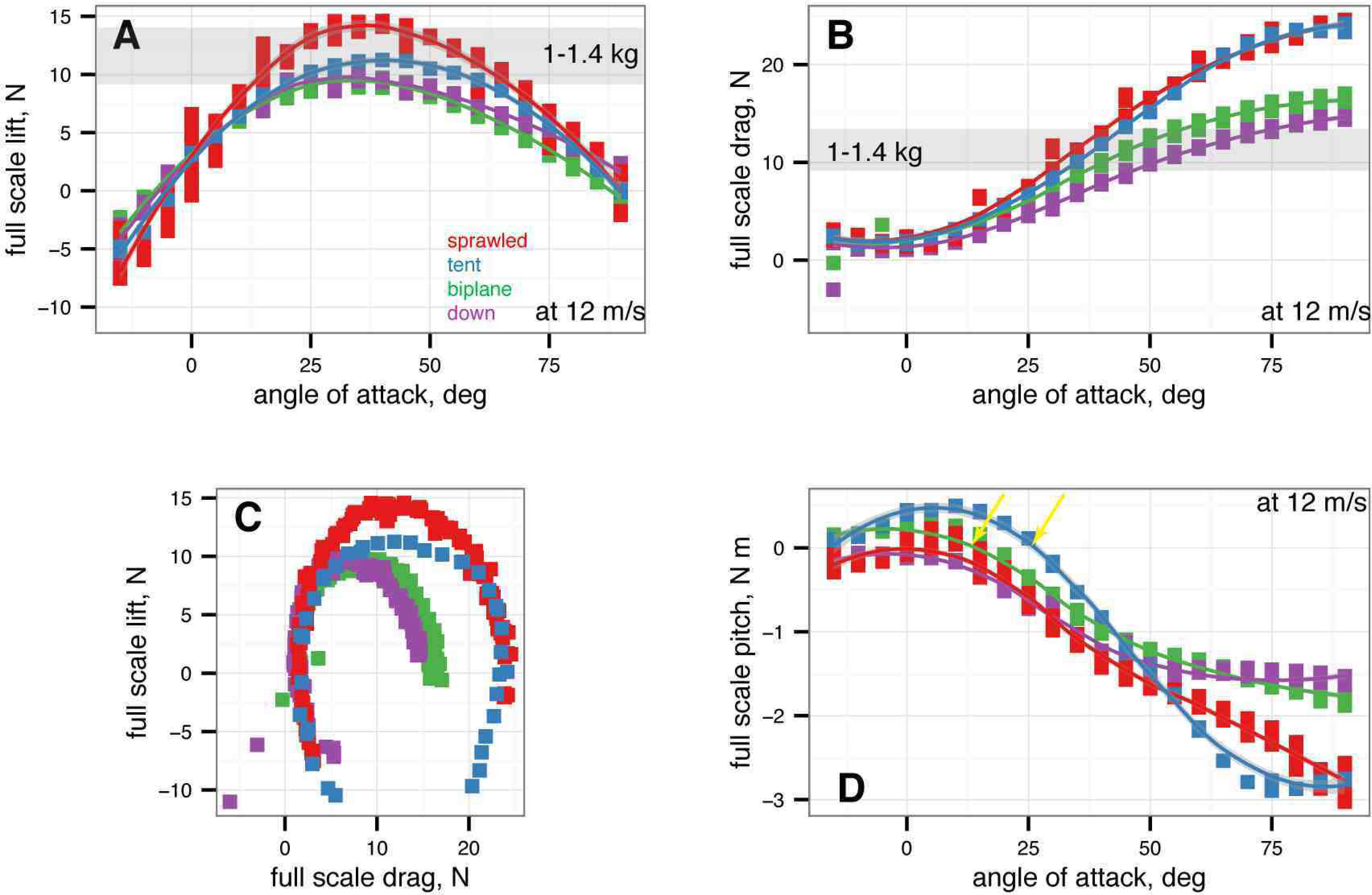
Symmetric wing pronation/supination control effectiveness for tent posture for wing angles of –30° (large down triangle), –15° (down triangle), 0°(square), +15° (up triangle) and + 30° (large up triangle). Wing pronation/supination (wing angle of attack) is effective at changing the lift generated but exhibits reversal at high angle of attack where stall occurs.

### Control effectiveness of asymmetric wing positions

Figs. 18 through 20 give the control effectiveness for asymmetric wing movements, including asymmetric wing sweep (Fig. 18), asymmetric wing pronation (Fig. 19), and asymmetric wing tucking (Fig. 20). For asymmetric wing sweep, the wings are swept in opposite directions up to 45°. For asymmetric wing pronation, the wings are pitched in opposite directions (e.g. left wing up, right wing down) up to 30°. For asymmetric wing tucking, one wing is tucked in entirely.

**Figure 18.**
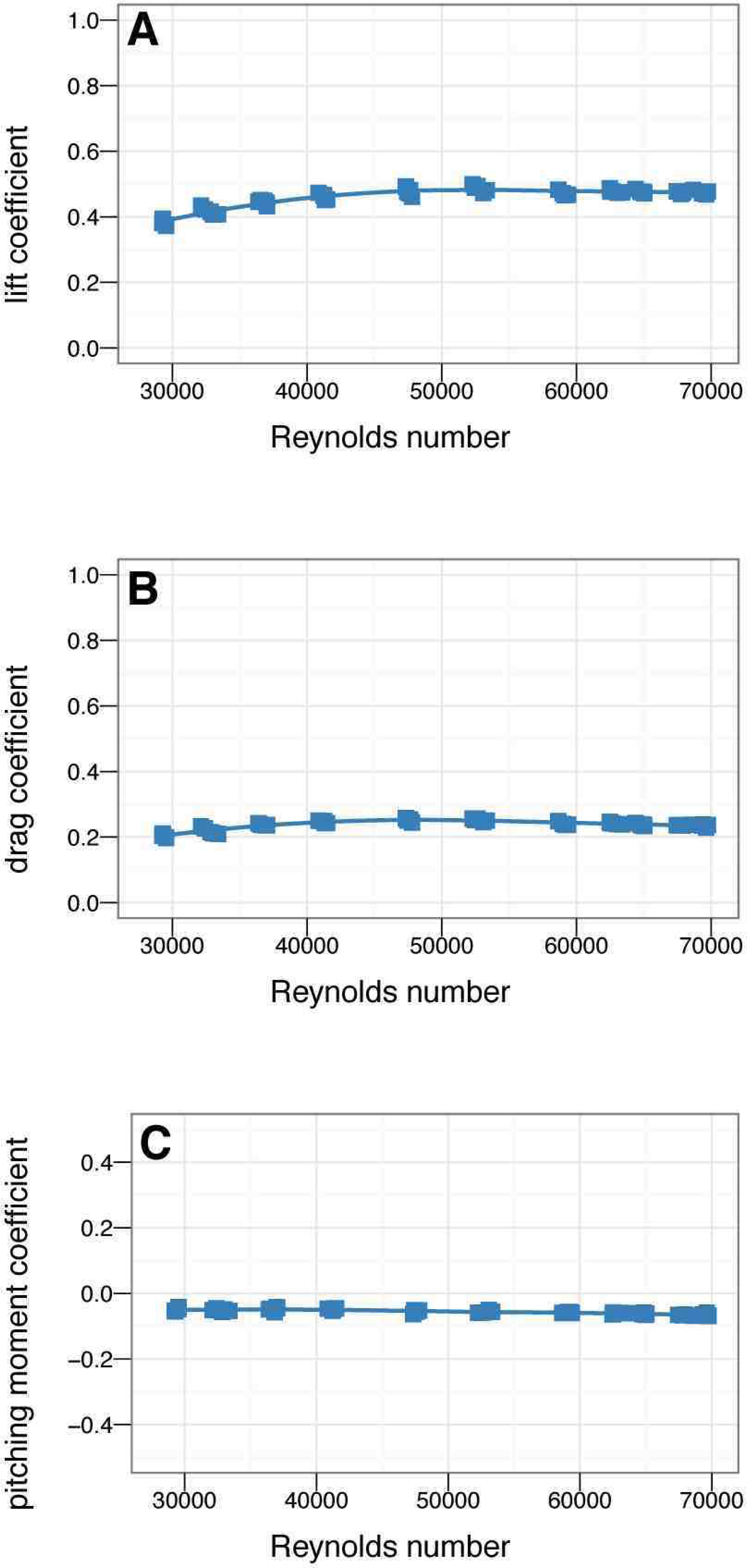
Asymmetric wing sweep (e.g. left and right wings swept forward and backward) control effectiveness for tent posture for wing sweep angles of –45° (large down triangle), –22.5° (down triangle), 0° (square), +22.5° (up triangle) and + 45° (large up triangle). Forward sweep generates upward pitching moments, backward sweep generates downward pitching moments. Considerable roll moments are also generated at higher angles of attack. Non-zero roll moments for symmetrical postures (B, squares) is due to slight sting misalignment during test, illustrating the measurement noise of the test.

**Figure 19.**
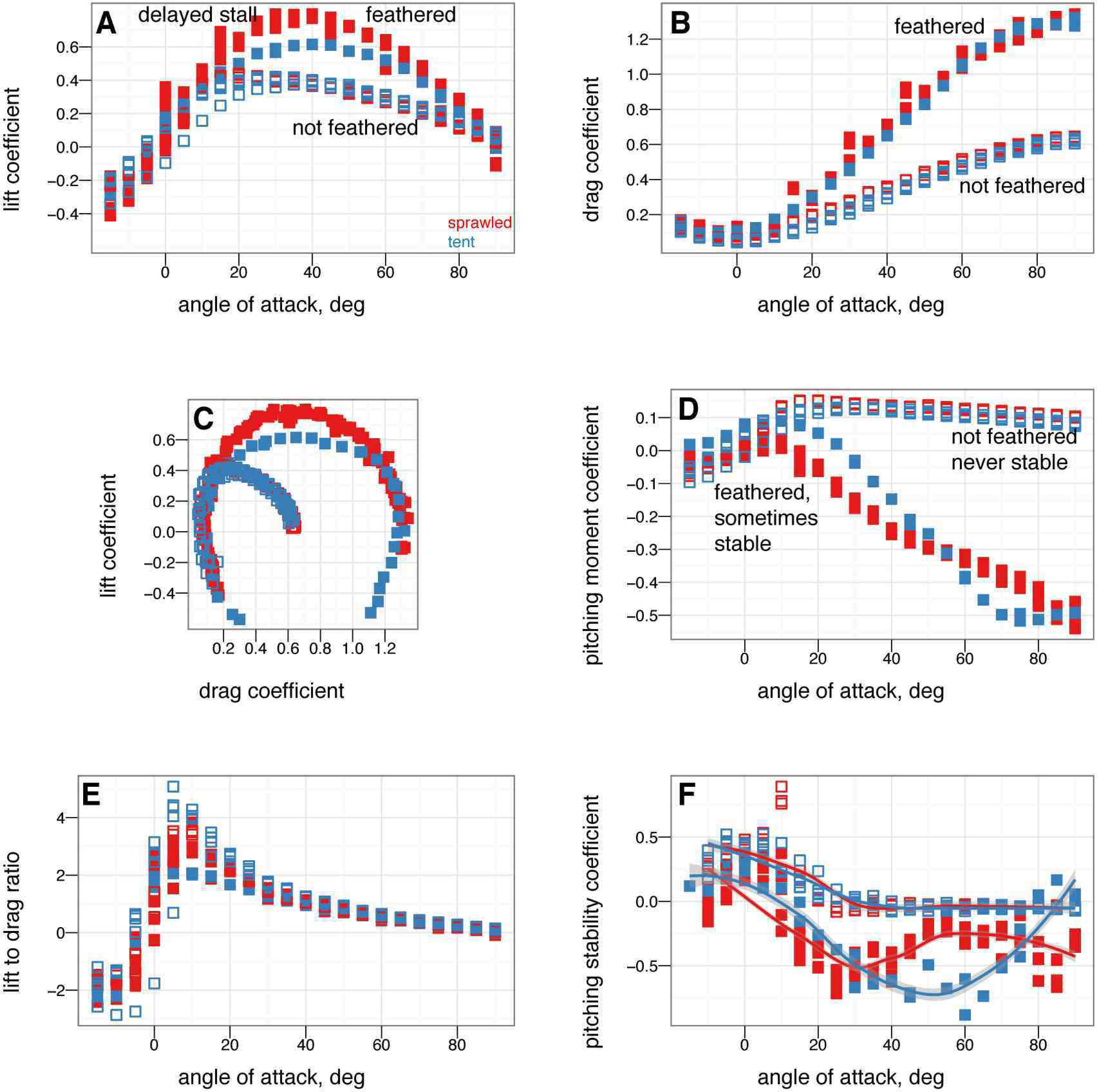
Asymmetric wing pronation (e.g. left and right wings pitched in opposite directions) control effectiveness for tent posture for wing pronation angles of –30° (large down triangle), –15° (down triangle), 0° (square), +15° (up triangle) and + 30° (large up triangle). At low angles of attack, asymmetric wing pronation generates large rolling moments. At high angles of attack, there is a shift in function and asymmetric wing pronation tends to generate yawing moments instead of rolling moments. Function at high angle of attack is similar to what is observed in human skydivers [37, 38]. Organisms may have navigated this transition from high angle of attack to low.

**Figure 20.**
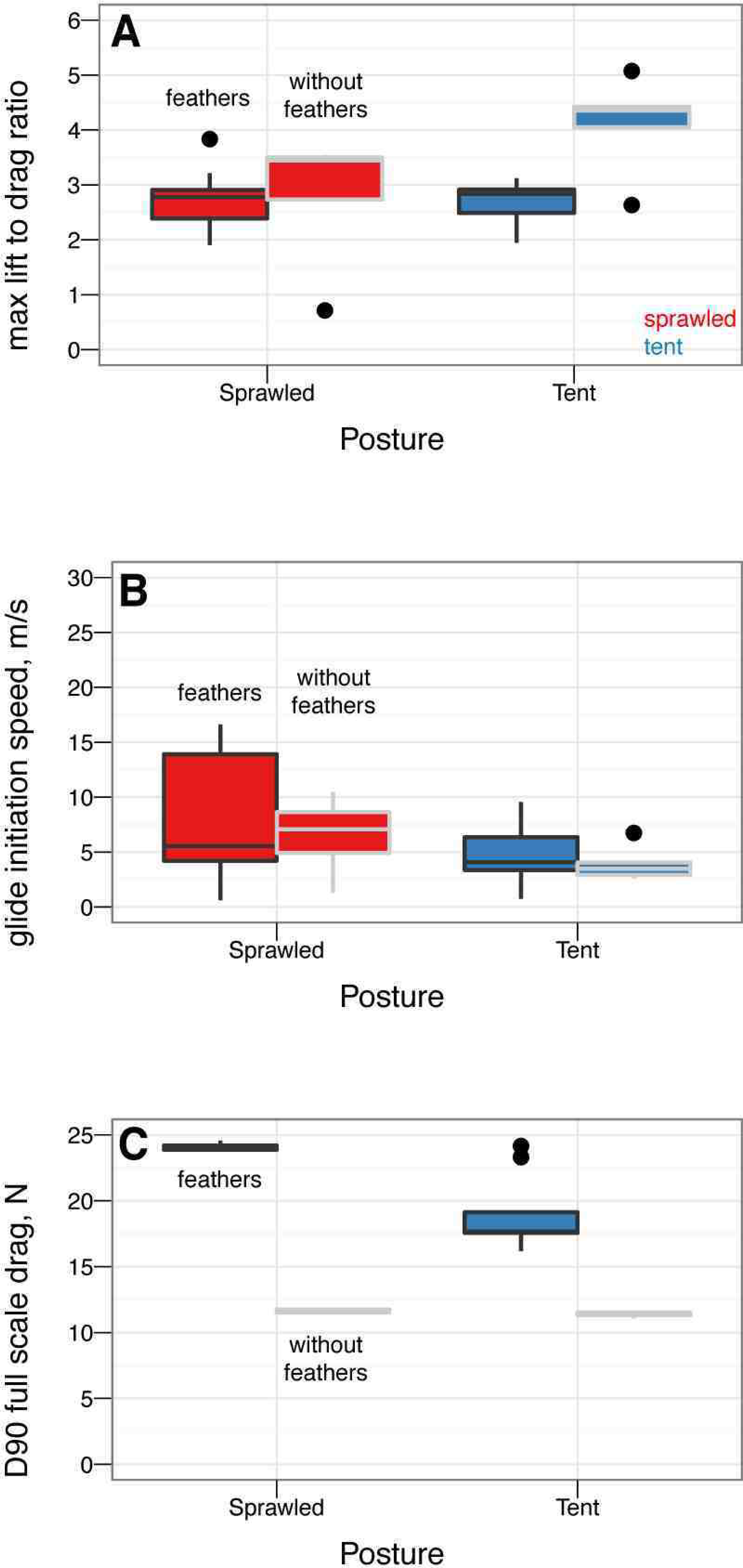
Asymmetric wing tucking control effectiveness for tent posture; both wings out (solid square), no right wing (open square) and no wings (open diamond). Tucking one wing produces large roll moments but at the expense of one quarter of the lift. Large yaw moments are not generated except at higher angles of attack where the leg and tail positions become more important. Rolling moments generated in the two-wing symmetric position illustrates the senstivity of symmetry, model positioning, and sting placement; in addition, yawing moments at extreme angle of attack further illustrate sensitivity to position which could be exploited as a control mechanism during high angle of attack flight.

### Control effectiveness of asymmetric leg positions in yaw

Control effectiveness of asymmetric leg positions in yaw is plotted in Fig. 21 and S3–S5 (available online).

**Figure 21.**
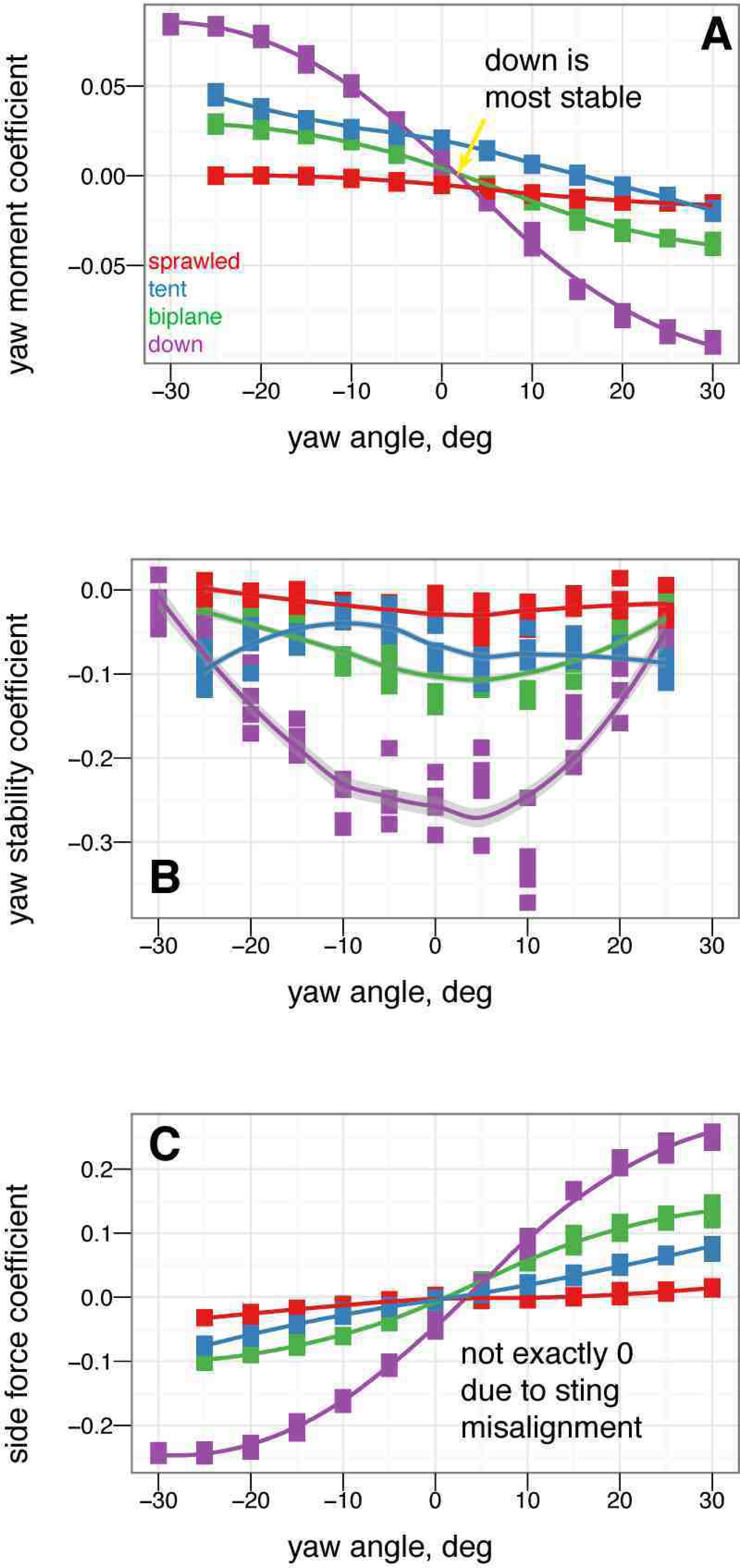
Asymmetric leg dihedral (leg *dégagé*, see inset) effect on yaw. Baseline down position (solid square) versus one leg at 45° dihedral (down arrow). Placing one leg at a dihedral is destabilizing in yaw and produces side force and rolling and yawing moments due to the asymmetry.

### Control effectiveness of other asymmetric positions in yaw

The control effectiveness of some additional asymmetric tail and leg movements in yaw is given in Figs. 22–23 and S6 (available online), including lateral bending of the tail (Figs. 22 and S6) and placing one wing down (Fig. 23).

**Figure 22.**
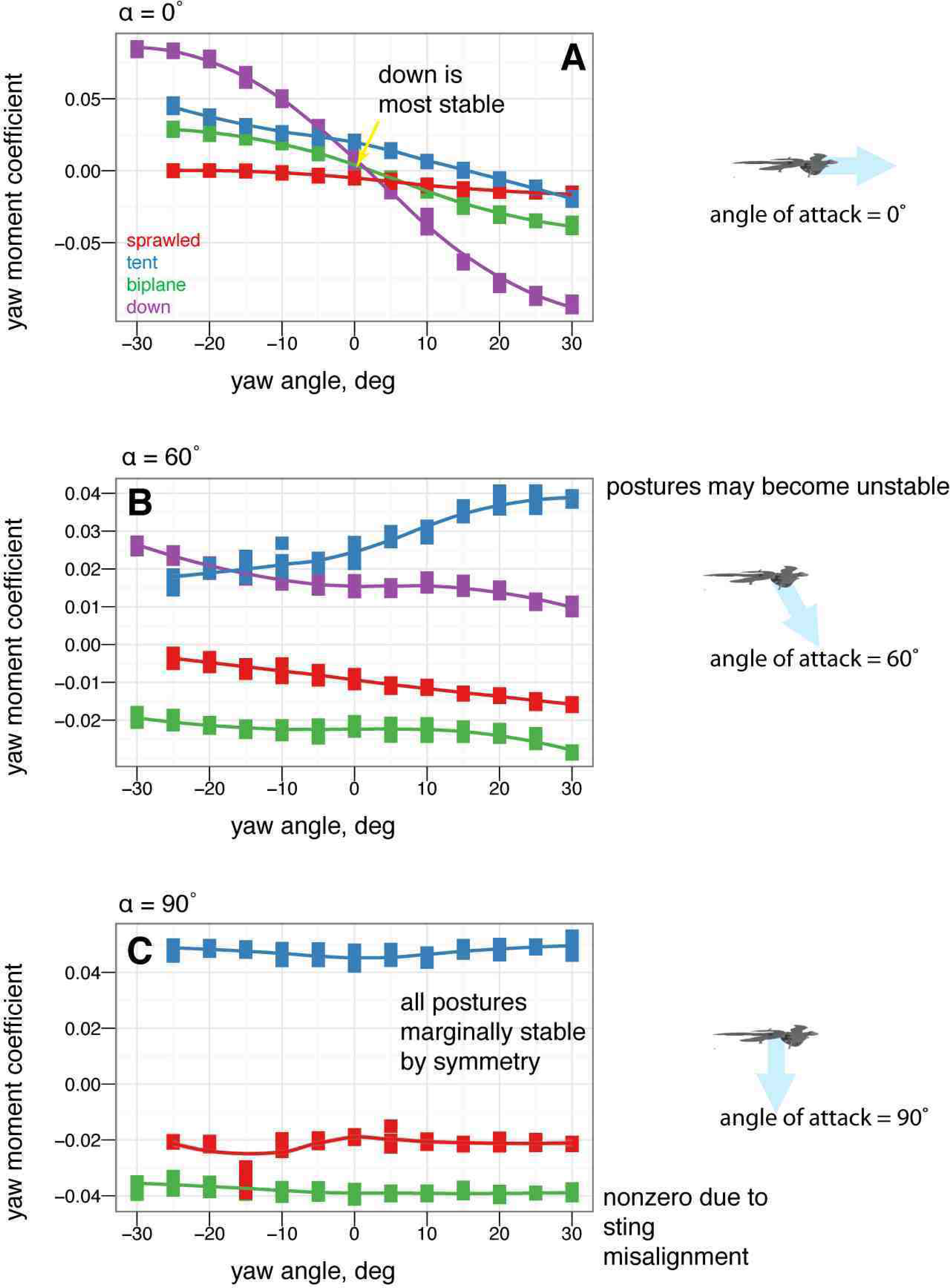
Asymmetric tail movement (lateral bending) effect on yaw, tent posture. Baseline tent position (solid square), tail 10° left (open square), tail 20° left (open triangle), tail 30° left (open diamond). The tail is effective at creating yawing moments but at low angles of attack it is shadowed by the body and larger movements are needed (yellow versus red lines).

**Figure 23.**
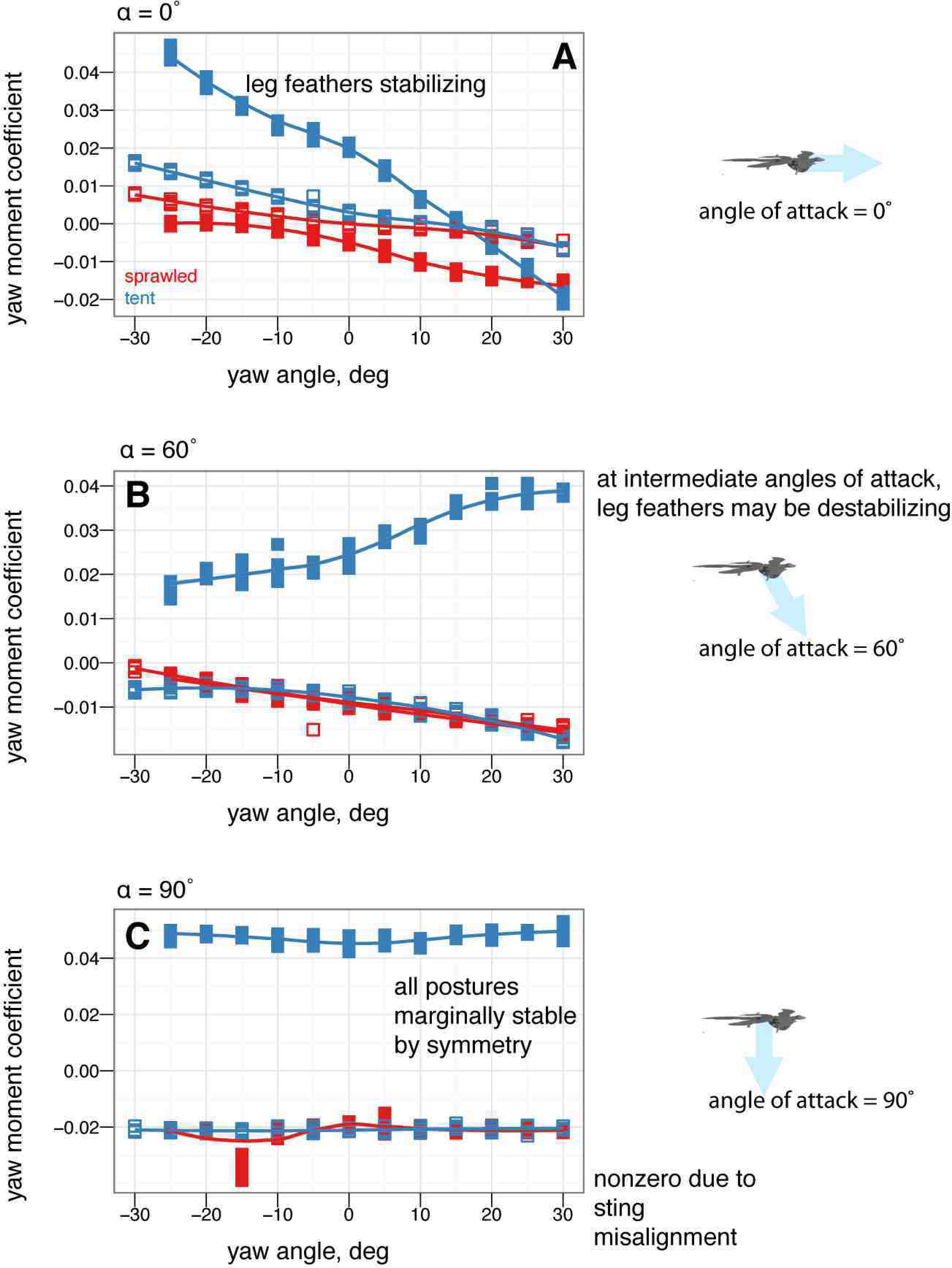
Asymmetric one wing down effect on yaw, tent posture. Baseline tent position (solid square), left wing down (down triangle). Placing one wing down does not make large yawing moments. Some roll and side force is produced at low angles of attack, at the expense of one quarter to one half of the lift.

## Discussion

### Postures have similar lift and drag coefficients but exhibit very different pitch (longitudinal) stability

All postures have roughly similar lift coefficients at low angles of attack (Fig. 2A); at high angles of attack, the main differences are due to the orientation and projected area of the legs. Baseline drag coefficients at zero lift (Fig. 2B) are similar to results measured in [57] (Dyke et al. Figure 1) within the scatter of the measurement, as well as to results for diving passerines in [67].

Examining the pitching moments reveals that only the biplane and tent postures have stable points (Fig. 2D). For the tent position, the stable glide angle is 35°, at roughly 12 m s^−1^ and an angle of attack of 27°. Xu et al. [34] also found the tent posture to be stable, which agrees with our results. For the biplane position, a stable equilibrium point appears at angle of attack 16°. The baseline sprawled posture, which possesses roughly equal fore and aft area, is marginally stable in pitch (in effect, the longitudinal center of pressure is at the center of mass), while the down posture is never stable because the legs are not employed in lift generation (the longitudinal center of pressure is ahead of the center of mass). For the sprawled posture, tail movement can be used to trim the body to longitudinal stability (Fig. 12); down posture can be trimmed to marginal stability using the tail (Fig. 11).

Anatomical criticisms [60] aside, for biplane postures, these stability results agree with [32], who argued from simulation results (that were highly dependent on parameter selection) that the biplane posture was stable. [57] also found this posture to be stable. In contrast, Xu et al. (as described on television in [34]), found the biplane to be unstable in wind tunnel tests except at high angle of attack. Alexander et al. [33] found that with nose-heavy ballasting, a sprawled/biplane posture could be made stable; we agree with this, with the caveat that such ballasting may not be biologically realistic as the densities of biological tissues do not vary as greatly as the density difference between lead and styrofoam.

Our predicted equilibrium glide angle for the tent position seems reasonable [50]. The animal would be fast enough to require some kind of landing maneuver to avoid injury [50]; using a simulation approach similar to [57, 68], one could evaluate the perching or landing ability of this animal using our data. Our glide angle and speed are higher than in [33], however, the weight estimate of Alexander et al. is half ours.

Based on projected full scale forces (Fig. 3) and stability considerations, we calculated the steady-state glide speed and glide angle from *C_L_* and *C_D_*, and estimate the †*M. gui* could glide in the baseline tent position at around 12 m s^−1^. The baseline sprawled posture and baseline down posture are unstable in pitch. The baseline biplane position at 12m s^−1^ does not appear to generate sufficient resultant force (lift and drag) to support body weight (1–1.4 kg) estimated by scaling based on [69–71], also from [72]; see Methods) at a speed of 12 m s^−1^. †*M. gui* would have had to move through the air at faster speeds to generate enough aerodynamic force to balance its weight. We did not mechanically evaluate if feathers cantilevered out the feet in the style of muffed feet on pigeons is able to carry significant loads; however this was a common point of mechanical failure in our physical models, suggesting it would have been a limitation for that hypothetical posture.

At first glance, there also appear to be differences in the maximum lift to drag ratio, minimum glide initiation speed, and parachuting drag for different postures (Supporting Figs. S1 and S2). It is important to note that these “optima” (maximum *L/D* optimizes steady glide distance; maximum parachuting drag optimizes straight-down fall velocity) reflect a very narrow criteria of optimality and are not always achievable because of constraints, such as from stability or anatomy. In particular, none of the most optimal configurations are stable. Blind application of gross aerodynamic performance parameters (such as [65]) may be misleading if it ignores other constraints.

### Coefficients are insensitive to Reynolds number

The Reynolds number sweep (Fig. 4, Table 1) shows that the models under test here are in a regime where aerodynamic coefficients are relatively insensitive to Reynolds number, so that results are valid for the full-scale †*M. gui*, as well as for full-scale †*Archaeopteryx*. This result was briefly discussed in [50] but additional details are relevant here. Unlike in gliding ants [36] or in typical low Reynolds number structures such as crab antennules [73] or blastoid respiratory hydrospires [74], there are not shifts in function of the wings as Reynolds number is varied over a range of sizes and speeds (Fig. 4). This is similar to what is observed in wind tunnel models of *Draco* lizards (Evangelista, in preparation) and Anna’s Hummingbirds (*Calypte anna*) (Evangelista, in preparation) and is similar to what is expected from typical high Reynolds number aerodynamics [50, 75–77]. In aerodynamic model tests of engineering airfoil sections with tripping, similar results are seen in this Reynolds regime [78] or in “rough” wings [79]. The absence of scale effects here provides added assurance that these results should be broadly applicable in evaluating maneuvering during evolution or ontogeny.

### Leg and tail feathers have important implications for aerodynamics and stability

Leg feathers forming a hindwing will stall at higher angles of attack than without a forelimb wing ahead of them (Fig. 5A, similar to a jib and a mainsail, or flaps on an airplane; alternatively, tandem wings have similar effects). Leg feathers also increased drag at high angles of attack (Fig. 5B) and altered stability (Fig. 5D). None of the shapes tested were stable without leg feathers present (Fig. 5D). This suggests that leg and tail morphology in fossils may be informative as to the stable glide angles or postures an organism can adopt in the air. The leg feathers were initially downplayed as a taphonomic artifact [80]; however subsequent finds of a wealth of specimens with feathers on the legs and tail [5,7,8,28] beg further work to evaluate their aerodynamic significance in a comparative framework.

Leg feathers increased *D*_90_ and decreased the lift to drag ratio, however, without leg feathers the models were not stable (Fig. 6). Higher *L/D* without leg feathers may be achieved by reduced drag from surfaces whose ability to produce lift is limited by their downstream location behind the forewings. This may have promoted an evolutionary shift from body forms with feathered legs form to forms with large forewings and reduced legs (as is seen in the evolution of birds) [39, 56].

The stability afforded to some postures by leg feathers is important to consider. For example, considering *L/D* ratios alone, †*M. gui* in tent position with no leg feathers might be expected to glide at speeds 1.4× faster (about 17 m s^−1^) compared to the baseline with feathers. However, the stability results show that without closed-loop control, an †*M. gui* without leg feathers would pitch upwards until stalling, and then tumble. This illustrates once again that assuming “better glide performance” is a single number such as *L/D* is an oversimplification; higher *L/D* means higher long distance glide performance only, and only when stability or control enables it to fly such trajectories. High *L/D* does not mean lower glide speed. Furthermore, long distance glide performance may not be the only performance task of interest, especially in a constrained or cluttered environment like a forest. For comparison, among human skydivers, steep approaches are often used to build speed in order to enable finer control near the ground. This is also the logic behind steep final approaches in powered aircraft, as it reduces the impact on control of an engine failure near the ground; and in the precision landing event during competition skydiving.

Living animals differ from models in being dynamic and that the various postures evaluated in this study (and others) might have been used in different circumstances to maximize the aerodynamic potential of the living animal. Dynamic behaviors (flapping, inertial flailing) could increase the maneuvering abilities further beyond what is discussed here, but these results provide a useful first-order understanding.

### Yaw stability depends on posture and leg feathers, and exhibits shifts based on angle of attack

Stability varies in different axes (pitch, versus yaw and roll). A particular shape and orientation relative to the flow which is stable in pitch may not be stable in the other axes.

Some postures (notably legs-down) were observed to be more stable than others in yaw (Fig. 7). More importantly, postures which are stable at low angle of attack (such as tent) were unstable at intermediate angle of attack, and all postures were marginally stable at 90° angle of attack (Fig. 8). Leg feathers were similarly seen to have different effects on stability with angle of attack (Fig. 9). The significance of this result is that during a shift from high angle of attack directed aerial descent, through mid-angle of attack gliding, to low-angle of attack flight, different plan forms have different stability characteristics in yaw. The aerodynamic basis for the difference is not yet clear, although it is likely due to effects of vortex shedding or separation at the tips and trailing edges of the various aerodynamic surfaces or the body itself (such as the stabilizing mechanism for high angle of attack lifting bodies). While some like to artificially divide parachuting and gliding from “true” (flapping) flight, both can be more dynamic and unsteady than the terms often imply to the casual reader, as seen here even in static stability and control effectiveness. Further work is needed to examine the basis for the shifts, using flow visualization, and to consider aerial behaviors as a continuum of maneuvering ability [1].

### Control effectiveness varies with angle of attack and can exhibit reversal or shifts from one axis to another

Control effectiveness was observed to vary with angle of attack (Fig. 8, 9; Fig. 10 onwards). Furthermore, there were cases in which its sign completely switched, i.e. when a control surface does the opposite of what it normally does (Fig. 11, down posture with the tail in pitch; Fig. 13, tent posture with the tail in pitch; Fig. 17, wing pronation in tent posture). These happen in pitch at high angles of attack and in yaw at different angles of attack and postures. Reversal during abnormal operating conditions in aircraft and ships can cause collisions and crashes. In a biological system, the examples of reversal here represent complete shifts in the function of an appendage that would happen coincident with a transition (evolutionary or during a maneuver) from steep-angle directed aerial descent to lower angle of attack aerial behaviors. This deserves further study; the basis for reversal is unclear in these models and flow visualization is needed.

As with the other measurements, removal of leg feathers tended to eliminate control effectiveness (for example, Fig. 12C versus D). This might suggest that as birds evolved and moved away from long tails and feathered legs, the control effectiveness that those surfaces once possessed became reduced, or possibly was shifted to another surface (the forelimbs/wings). This is bolstered by the observation that birds with partially amputated tails (such as caused by attacks by household cats) can still fly. In the data presented here, wing sweep (in a manner similar to steering a windsurfing rig) was very effective at creating pitching moments, similar to patterns seen in model tests of diving Anna’s Hummingbirds (*Calypte anna*) (Evangelista, in preparation). Forward sweep also appeared to increase the maximum lift coefficient, which could allow slower flight speeds; wing pronation had similar effects.

Further comparative study of wings, tails and putative empennage in general, including reference to convergent examples in pterosaurs, are discussed in [56] and in chapter 3 of [39], and other later work [81].

### Some asymmetric movements are effective in rolling or yawing

For asymmetric wing movements, similar trends were observed. Asymmetric wing sweep was effective (Fig. 18). Tucking one wing (Fig. 20) was effective in rolling. Other work has observed use of this particular movement in rolling maneuvers in young birds [39]. Asymmetric wing pronation, in particular, tent posture with one wing changing its pronation/supination, was observed to produce large rolling moments at low angle of attack but large yawing moments at high angle of attack (Fig. 19). The function of such motion in creating yaw at high angle of attack is similar to certain arm positions used in human skydiving to create yaws [37, 38]. In the context here, this is another observation of a major shift in the function of a control surface with angle of attack. Our results demonstrate that as an organism transitions from high angle of attack directed aerial descent to lower angle of attack aerial behaviors, the function of the wings in control changes.

On the other hand, certain asymmetric movements such as placing a leg *dégagé* (Fig. 21) or *arabesque* (Figs. S4 and S5) or placing one wing down to attempt to create yawing moemnts (Fig. 23) had surprisingly little effect on yaw, roll, or side force, and also had the negative consequence of the loss of a large portion of lift. There would have been little selective advantage for using these asymmetric postures given that there are more effective means of producing yaws, rolls, and side forces.

Asymmetric tail movements (lateral bending) were only partly effective compared to forelimb wing movements (Figs. 22–23, S6). At low angles of attack, the tail may be shadowed by the body, e.g. it is downwind of the body and because of body-tail interactions, has little flow, which result in reduced control effectiveness. As an organism’s flight environment shifts from high angle of attack directed aerial descent to low angle of attack aerial behaviors, surfaces that were effective at high angle of attack may become less effective due to these effects.

### Possibility for animals to alter their trim and stable point?

While the baseline sprawled, down, and biplane postures were largely unstable, the control effectiveness sweeps show that some degree of trim control (alteration of the stable point by altering wing sweep, tail angle, or some other movement with large enough control effectiveness) may have been possible to help maintain those postures. This is done by soaring birds in order to reduce speed and fly slowly at minimum sink speeds while thermalling. Thermalling is a very derived behavior, but we have every reason to expect animals to use all available control channels even early during the evolution of flight. Due to the factorial growth in runs required to explore multiple permutations of posture and multiple combined appendage movements, it was not possible to fully explore such combination effects (and, indeed, when considering closed loop control, it may not be worthwhile to delve too deeply into such a series). In other work, we have observed such shifts in the stable point, for example, in human skydivers [37, 38] and during dive pullout in Anna’s Hummingbirds (Evangelista, in preparation). More importantly, we have identified several control channels that are effective (e.g. symmetric wing sweep, asymmetric wing pronation, tail movement), as well as many that are not effective in comparison.

### Tradeoff between stability and maneuverability

The agility of an animal is the combined result of both stability and control effectiveness [10,11,30]. Many have proposed a trade-off between aerodynamic stability and maneuverability [10,54,82–85]; a stable form is “easier” to control but slow to respond, while an unstable form would require high control effectiveness and good sensorimotor control but could potentially respond more quickly. Unfortunately, data from previous studies does not provide strong evidence for such a tradeoff. Past model studies of gliding frogs computed stability indices using a noisy sensor and setup and 20° angle increments [10, 11, 30], and were thus not as accurate as those reported here. Other studies of gliding frog stability provided only qualitative assement of stability, recording whether frog modes of different morphologies and postures tumbled or not [54].

It is difficult to draw stronger conclusions about stability-maneuverability tradeoffs solely from the data in this paper. Simple glide metrics (after [54], which [10] considers also as metrics of bank turns, e.g. *L* cos *ϕ* and *L* sin *ϕ* with *ϕ* arbitrarily fixed at 60°) show few significant differences (Figs. S1 and S2 are probably not informative in this respect). Removal of leg and tail feathers reduced stability but also removed the control effectiveness of those surfaces. On the other hand, there are large differences in stability for different postures, different angles of attack, and different glide angles, including roll and yaw, as well as differences in which movements are effective and which are not. This suggests our measurements may be informative to consider in understanding how sensorimotor and flight control abilities (which do not fossilize and cannot be observed directly) may have changed during evolution.

### Maneuvering must be considered when considering the evolution of flight in vertebrates

Taken together, these results show that morphology can have large effects on the stability and control effectiveness. Stability and control also place constraints on aerodynamic performance (specifically, whether or not reduced glide angles, lower glide speeds, or improved parachuting performance can actually be achieved). It is clear that even animals with little obvious aerial adaptations posess some degree of stability and control [10, 36–38, 40]. Stability and control effectiveness of appendages would have changed as bodies and appendages changed, and also as the flight regime changed from one of steep glide angles and angles of attack, as might occur during directed aerial descent [1] early the evolution of flight, to one of shallower glide angles and lower angles of attack. Observations of *L/D* and more traditional glide performance from simulations also support this [57]. The changes in tail and leg morphology during the transition from theropods to birds (and convergent changes from early pterosaurs to later pterosaurs and early bats to later bats [13]) beg for the metrics observed here to be studied in a phylogenetic comparative context [39, 56], to examine how they change as the morphologies are changed and to examine what skeletal or other features co-occur with changes in aerodynamics; additional study is also needed dynamics of high angle of attack maneuvers [36, 39, 86] and responses to aerial perturbations [36, 40, 87].

## Materials and Methods

### Models and postures

Scale models of †*M. gui* (scale model snout-vent length 8 cm were constructed from published reconstructions and from photographs of the fossils [2, 32, 34, 58]. The models are shown in Fig. 1A. Model construction was guided by dissection of Starlings (*Sturnus vulgaris)*, reference to preserved specimens of birds, bird wings, and lizards, casts of †*Archaeopteryx*, and illustrations in textbooks on vertebrate functional morphology and vertebrate paleontology [13, 88]. Photographs of the †*M. gui* holotype IVPP V13352 were printed on a laser printer (Xerox, Norwalk, CT) at full scale and at model scale to further guide model construction.

Models were built on an aluminum plate with polymer clay (Polyform Products Co., Elk Grove, IL) to fill out the body using methods described in [50]. Removable tails and heads, to allow repositioning, were constructed using polymer clay over steel rods. The forelimbs were constructed by bending 26-gauge steel wire scaled to the lengths of the humerus, radius and ulna, and digits as seen in published photographs of the holotype. Similarly, hindlimbs were constructed with wire scaled to the lengths of the femur, tibiotarsus, tarsometatarsus, and digits. For the appendages and tail, feathered surfaces were modeled using paper and surgical tape (3M, St. Paul, MN) stiffened by addition of monofilament line at the locations of the individual feather rachises. Tape was also used on the leading edge of all surfaces trip the boundary layer into turbulence [78, 79]. This method of creating wing surfaces was compared to wings with craft feathers individually sewn onto them and seen to provide equivalent results [50]. In addition, models of Anna’s Hummingbirds (*Calypte anna*) constructed using the same techniques have been shown to faithfully reproduce the aerodynamic properties of diving hummingbirds (Evangelista, in preparation).

The postures of the models (Fig. 1B-E) were chosen based on those previously published [2, 32,34,58]; others [59, 81] were not yet proposed at the time experiments were done. Some of these postures are anatomically dubious. We recognize that some of the postures tested are less feasible than others. The approach taken here is to test all previously proposed reconstructions in order to examine the aerodynamic implications of these shapes from a purely physical standpoint. In particular the sprawled posture drawn in [2] has been criticized, because interference between the trochanter on the femur and the surrounding structures of the ischium should have made that posture difficult to assume [13, 34, 88]. However, Xu never intended the sprawled posture as an actual reconstruction *per se* but rather just a convenient way to illustrate the planform [89]. In the absence of fossil material illustrating otherwise there is generally no reason to assume extraordinary hip anatomy not seen in any other theropod. Similarly, a feasible mechanism for maintaining feathers in the biplane / muffed feet posture of [32] under load has never been proposed, with some authors questioning the position entirely [60] and others supporting it [33, 57]. We also tested models in postures more strongly inferred for theropods, including a legs-down posture with leg abduction limited to ≤ 45° [34], and a tent posture in which the legs are extended caudad with the feathered surface extending over the proximal part of the tail [34, 58].

With the uncertainties inherent in applying a physical modeling approach to an extinct animal with only a single published skeleton, statements about aerodynamic performance in †*M. gui* should always be taken with a grain of salt.

### Conditions for dynamic similarity and Reynolds number sweep

If a model and organism are dynamically similar, then the ratio of forces acting on corresponding elements of the fluid and the boundary surfaces in the two is constant, and force and moment measurements on the model can be scaled to calculate forces and moments acting on the organism [75]. To achieve dynamic similarity in model tests of aerodynamic maneuvering, the Reynolds number (Re = *uL/ν*) should match. Reynolds number (Re = *uL/ν*, where *u* is the velocity of the fluid, *L* is a linear dimension; snout-vent length in this study, and *ν* = 15 × 10^−6^ m^2^ s^−1^ is the kinematic viscosity of air) is the nondimensional ratio of viscous to inertial forces. Based on pilot studies we estimated Re for the full scale †*M. gui* to be approximately 200,000. Limitations on the wind tunnel size and speed required the Reynolds number of the model to be 32,000. Model tests at lower Reynolds number may be acceptable if it is possible to verify that scale effects are not present, and if the flow regime is the same between model and prototype.

Early in the evolution of animal flight, organisms likely flew at moderate speeds and high angles of attack where flows appear like bluff body turbulent flows (in which coefficients are largely independent of Re, for 10^3^ *<* Re *<* 10^6^). We performed a sweep of wind tunnel speed, to examine Re from 30,000 to 70,000, to validate that scale effects were not present. As additional support for this approach, tests for maneuvering bodies are nearly always tested at well below full scale Re, e.g. the largest US Navy freely-maneuvering model tests are well below ⅓-scale. Our methods were also benchmarked using model tests at full scale Re of *Draco* lizards and Anna’s Hummingbirds in glide and extreme dive pullout maneuvers compared to live animal data (Evangelista, in preparation).

### Force measurements

As described in [50], models were mounted on a six-axis force transducer (Nano17, ATI Industrial Automation, Apex, NC), which was in turn mounted on a 1/4–20 threaded rod damped with rubber tubing, and attached to a tripod head used to adjust angle of attack. The force sensor and sting exited the model on the right side of the body mid-torso at approximately the center of mass. As a major source of measurement uncertainty was the positioning and mounting of the model on the sting, models were repositioned and remounted for each replicate run.

Wind tunnel tests were conducted in an open jet wind tunnel with a 15 × 15 × 18 inch (38.1 × 38.1 cm) working section used previously for studies of gliding frogs [10, 30]. Tunnel speed was controlled using a variable autotransformer (PowerStat, Superior Electric Company, Bridgeport, CT) and monitored using a hot wire anemometer (Series 2440, Kurz Instrument Co., Monterey, CA).

As the wind tunnel dimensions are not as large as might be desired, windspeed profiles were taken which found speed across the tunnel width was within 2% of the mean at stations from 7.6–30.6 cm. This check suggests the shear effects should be negligible. In addition, as part of benchmarking before testing, smaller dinosaur models and models of other taxa were tested and found to have comparable force and moment coefficients to the final results and to results from tests in a larger wind tunnel. Tow tank tests of *Cephalotes* ants in which the ant is comparable to the size of the tank also have shown little effect on the moments [36].

Force transducer readings were recorded at 1000 Hz sampling frequency using a National Instruments 6251 data acquisition card (National Instruments, Austin, TX). Since the sensor was fixed to the model, the raw measurements were initially in a frame fixed to the model. Raw measurements were rotated to a frame aligned with the wind tunnel and flow using the combined roll, pitch, and yaw angles by multiplication with three Euler rotation matrices. Transformed measurements were then averaged over a one-minute recording. For each measurement, wind tunnel speed was recorded and used to compute Reynolds number. The sign convention for forces and moments is shown in Fig. 1.

Aerodynamics forces and moments were normalized to obtain nondimensional coefficients according to the following equations (using notation from [31]):

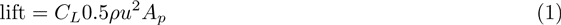

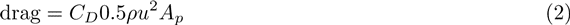

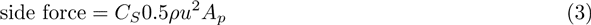

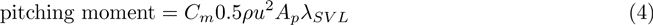

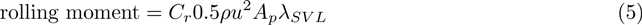

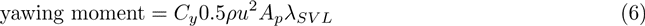

where *ρ* = 1.204 kg m^−3^ is the air density, *A_p_* is the model planform area, and *λ_SVL_* is the snout-vent length of the model. To allow comparisons among models, a single, consistent baseline configuration is needed. Accordingly, nondimensional coefficients are referenced to the planform area of the four-winged, sprawled position originally proposed in [2] unless specially noted. The questions of interest for this study are tied to the absolute value of forces and moments produced and differences that occur from the same animal in different postures; our choice of normalization preserves these distinctions in most cases. Coefficients were plotted in R [90] using the `ggplot2` library; all data were plotted with smoothed lines generated using loess smoothing only for visualization.

### Static stability coefficients

To assess static stability, we calculated nondimensional static stability coefficients from fixed-wing aircraft stability and control theory (notation from [31], see also [91–94]) and previously used in studies of gliding frogs [10, 30].

The pitching stability coefficient *C_m,α_* is defined as [10]

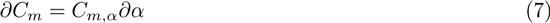

where *α* is the angle of attack and *C_m_* is the pitching moment coefficient as defined above. It is the local slope of the pitching moment curve, and is thus an indication of the sense (restoring if negative, or non-restoring if positive) and magnitude of the torque generated in response to a perturbation in angle of attack. If *C_m,α_ <* 0, the aerodynamic torque on the body will be opposite direction from that of the perturbation; this is the condition for static stability.

Similarly, for roll:

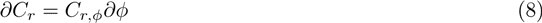

where *ϕ* is the roll angle and *C_r,φ_ <* 0 is the condition for static stability in roll. By symmetry, models at zero angle of attack have neutral rolling stability, and we did not calculate roll stability for most cases.

For yaw,

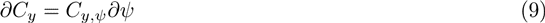

where *ψ* is the yaw angle and *C_y,ψ_ <* 0 is the condition for static stability in yaw (yaw stability is also known as directional stability).

Pitching stability coefficients were determined for models at different angles of attack (*α*), ranging from –15° to 90° at 5° increments. Yawing stability coefficients were obtained from models at different yaw angles (*ψ*) ranging from –30° to 30° at 10° increments. For each series of measurements, central differences were used to estimate the slopes at each point for each replicate run. Slopes were calculated from the measured coefficients using R [90].

### Control effectiveness of appendages

We also calculated nondimensional control effectiveness coefficients using methods from aeronautical engineering [92] used in previous studies of gliding frogs [30]. In general, control effectiveness for a control surface whose angular orientation relative to the flow can be changed is the partial derivative of the moment generated by the control surface with respect to the angle it is moved. High control effectiveness means a large moment is generated by a small movement of the control surface.

For tent and sprawled postures, control effectiveness was determined for symmetric and asymmetric movements of the feathered forelimbs (wings): symmetric protraction/retraction, asymmetric pronation, and complete tucking of one or both wings. Control effectiveness was also measured for feathered hind limbs/legs: asymmetric alteration of leg dihedral (for example, see Fig. 1H), lowering of a single leg, and change of leg relative angle to the body / angle of attack; and for the tail: dorsoventral and lateral bending. For these movements, we calculated the pitching control effectiveness as follows:

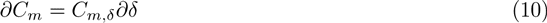

where *δ* is the angle of the control surface in question, with respect to neutral/baseline. Similarly, we calculated yawing control effectiveness for these surfaces as follows:

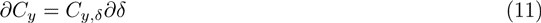

as well as rolling control effectiveness for asymmetric movements of the wings and legs:

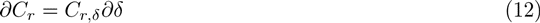

### Other flight performance metrics

To allow comparison with previous studies, two additional measures of maneuvering performance were computed: 1) the banked turn maneuvering index; and 2) the crabbed turn maneuvering index [10,30,65]. The banked turn maneuvering index assumes turns accomplished by banking is computed in two ways, both of which assume that some component of the lift generated is used to provide the force necessary for turning:

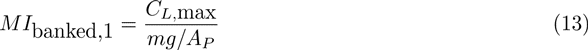

after [65] (note this is not a nondimensional index), and

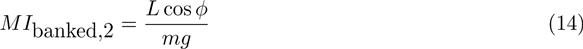

So that these indices could be compared to published values [10, 30], *ϕ* = 60° was used here, although the choice is arbitrary with no direct support from the fossils. Similarly, for crabbed turns, a nondimensional index is the horizontal component of side force normalized by body weight [10, 30]:

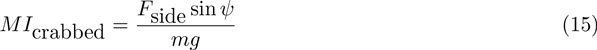

again with *ψ* = 60° arbitrarily chosen based on frogs [10, 30]. A valid criticism of these indices from [10, 30, 65] is that they are just scaled versions of *C_L_* and *C_S_* that are more informative, without having been manipulated by arbitrary choices of *ϕ* or *ψ*. These are included only for comparison to past literature.

Several flight performance metrics not immediately tied to maneuvering were also computed [10, 30, 54, 65]. As a measure of horizontal glide performance, we computed (*C_L_/C_D_*)_max_ for each posture [65]. Minimum glide speed, a measure of the ease of which gliding can be initiated, was also computed as *U*_min_ = [2*mg*/(*A_Pρ_C_L_*)]^1/2^ [65]. As a measure of parachuting ability of different postures, we also compared *D*_90_, the full scale drag for parachuting [65], as well as a nondimensionalized parachuting index *D*_90_/*mg*. [65] accepts a very limited definition of parachuting based on glide angle *<* 45°; however, gliding and parachuting are considerably more dynamic and unsteady than their names would imply and there are good reasons to consider aerial behaviors as a continuum of aerial maneuvering. These coefficients [65] are oversimplifications but are included here only for comparison to past literature.

### Estimation of mass and of location of the center of mass

The mass of a live †*M. gui* was estimated by scaling in two ways. One estimate was formed by scaling from published data for birds [69, 70] to estimate the mass and lengths of head, neck, wings, legs, body and then summing the masses and moments, methods identical to estimation of weights and centers for traditional naval architecture and other engineered systems. Another estimate was formed using scaling from many taxa based on long bone measurements [71]. Estimates of mass and of location of the center of mass fell within what has been published recently from a very detailed comparative study of archosaurs [72]. Masses (ranging from 1–1.4 kg, full scale snout-vent-length ∼ 35 cm) were used here only to estimate wing loadings and required glide speeds and to set the position of the sting.

## Acknowledgments

We thank five anonymous reviewers for helpful comments on the manuscript.

We thank the following undergraduate students, who also participated in the †*Microraptor* project over the years: Chang Chun, Michael Cohen, Vincent Howard, Shyam Jaini, Felicia Linn, Cyndi Lopez, Divya Manohara, Francis Wong, Karen Yang, Olivia Yu, and Richard Zhu. This research was done through the Berkeley Undergraduate Research Apprentice Program (URAP). We also thank Robert Dudley for occasional use of his wind tunnel, Tom Libby and the Berkeley Center for Integrative Biomechanics in Education and Research (CIBER) for use of a force sensor. We are sad to have lost one member of our team to tragedy, Alex Lowenstein, and we are grateful for our time with him.

This work was funded in part by an NSF Minority Graduate Research Fellowship (to DE), NSF Integrative Graduate Education and Research Traineeship (IGERT) #DGE-0903711 (to R Full, MK, R Dudley, and R Fearing), and the Virginia G and Robert E Gill Chair (to MK).

---

**Figure S1.**
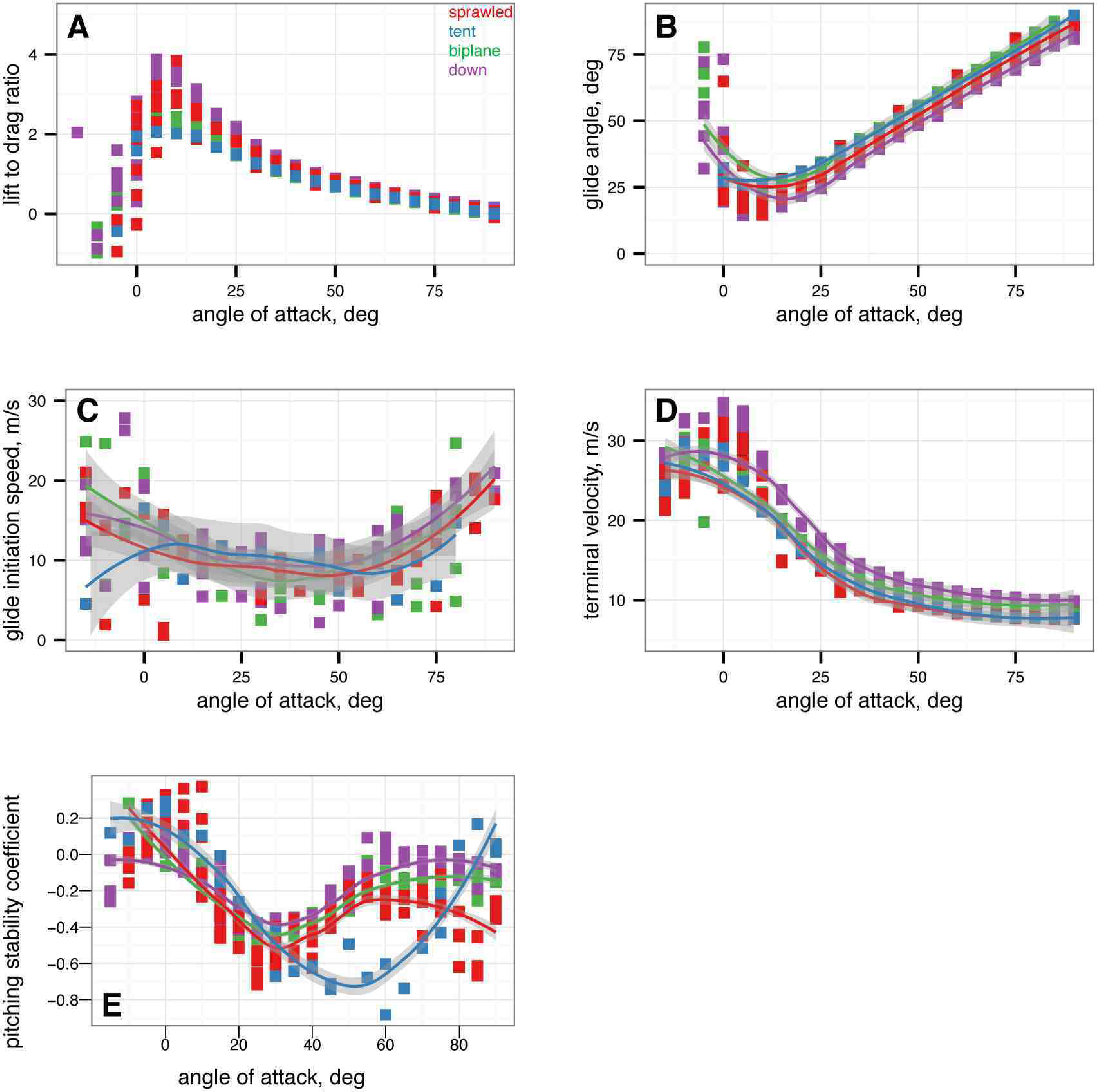
Simple gliding metrics after [65]. Red is sprawled, blue is tent, green is biplane, purple is down. *α* from −15° to 90° in 5° increments, with five or more replicates per treatment. A, Lift to drag ratio. B, Glide angle. C, Minimum glide speed. D, Terminal velocity (at which *D*_90_ = *mg*, assuming stability). e: Pitching stability coefficient (note pitching moment must also be zero for stable equilibrium).

**Figure S2.**
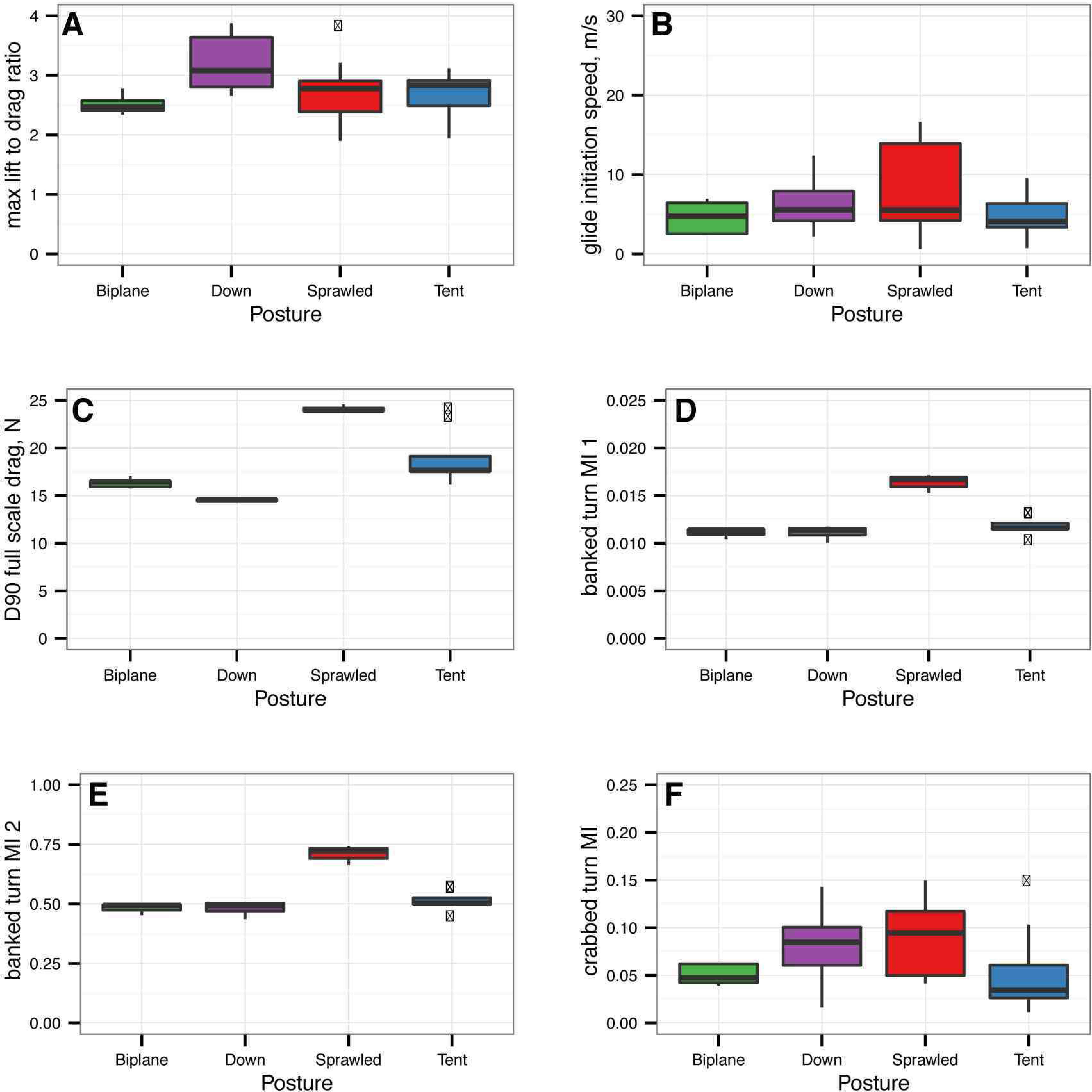
Comparison of simple glide metrics after [65] suggests the metrics are not informative. Red is sprawled, blue is tent, green is biplane, purple is down. A, Maximum lift to drag ratio, by posture, without regard to stability. [65]’s minimum ratio is never achieved because the models are not stable at the point where *L/D* is maximum. There is no difference in maximum lift to drag ratio among postures (Kruskal-Wallis, *P* = 0.1740). B, Minimum glide initiation speed, by posture, without regard to stability. The minimum speed is never achieved because the models are not stable at the point where *U_min_* is lowest. There is no difference in *U_min_* among postures (Kruskal-Wallis, *P* = 0.575). C, “Parachuting” drag, *D*_90_, by posture, without regard to stability. This drag is never achieved because the baseline postures are not stable at a 90° angle of attack. There are significant differences in *D*_90_ among postures (Kruskal-Wallis, *P* = 9.2 × 10^−5^); sprawled position has higher parachuting drag. D-E, banked turn maneuvering indices suggest sprawled posture may execute banked turns better than others, but posture is not stable. F, baseline postures not different in crabbed turn performance.

**Figure S3.**
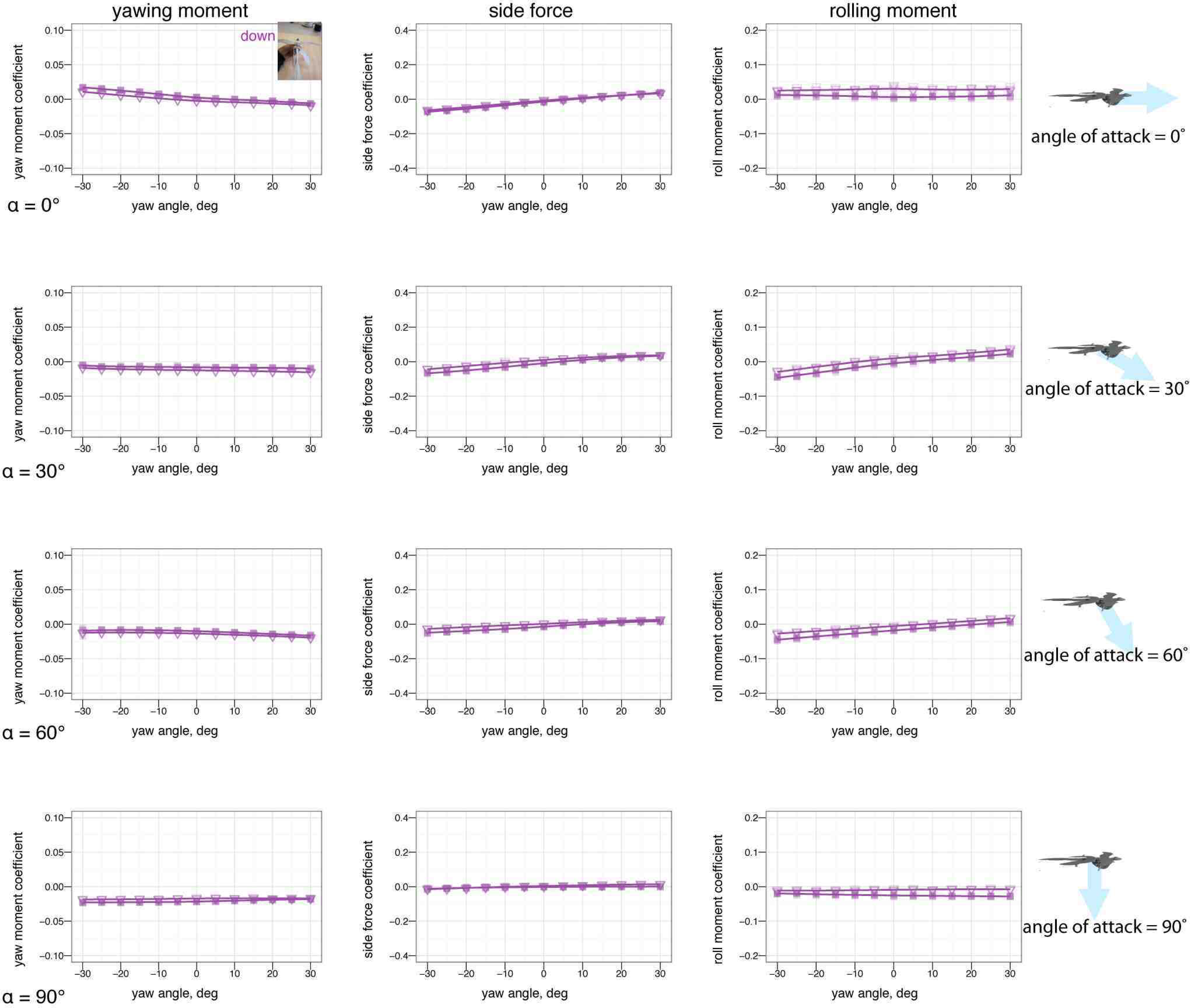
Asymmetric leg dihedral (leg *dégagé*, see inset) effect on yaw without leg or tail feathers. Baseline down position (solid square) versus one leg at 45° dihedral (down arrow). Without leg or tail feathers, the surfaces have little aerodynamic effect.

**Figure S4.**
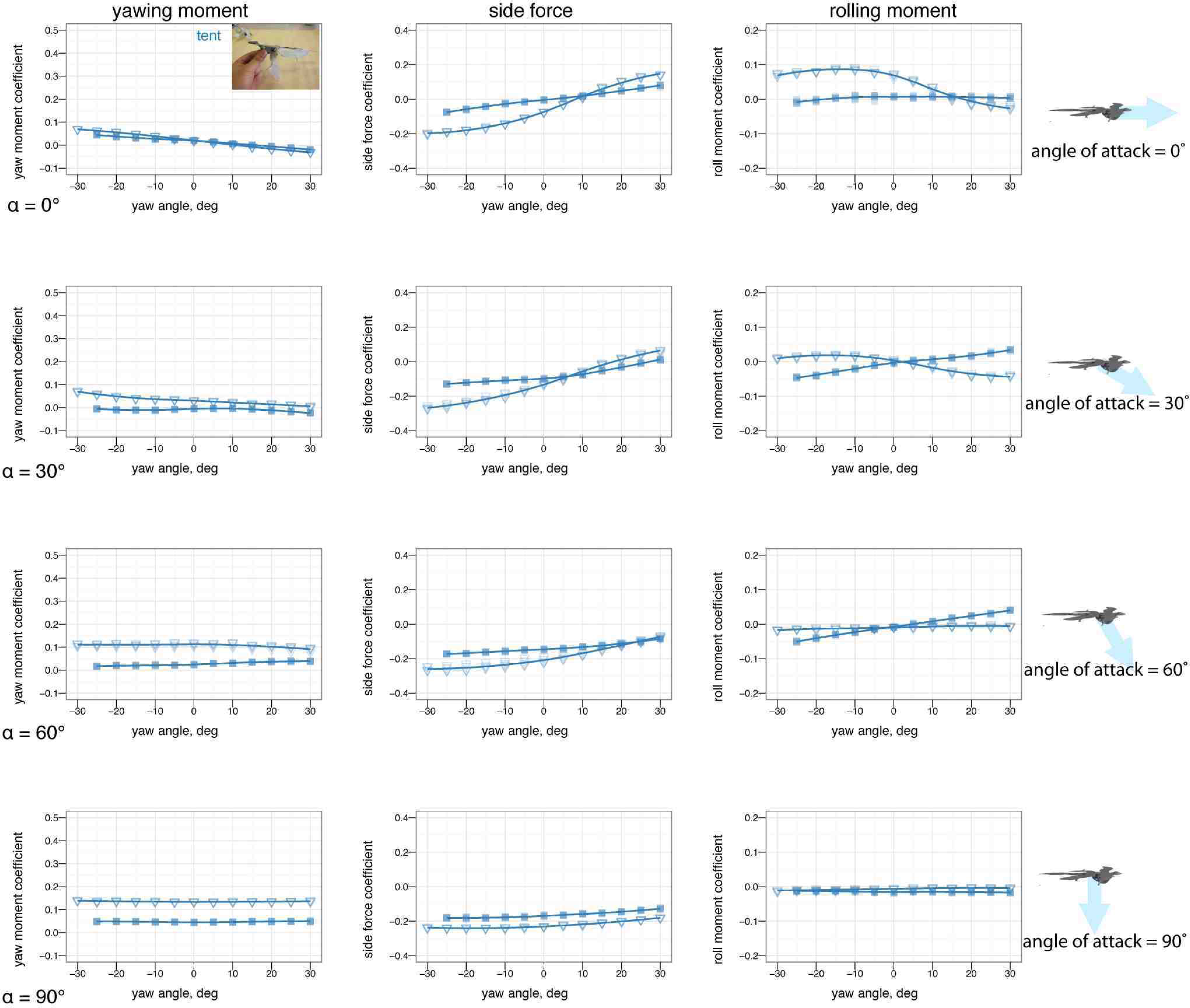
Asymmetric one leg down (leg *arabesque)* effect on yaw. Baseline tent position (solid square) versus one leg at 90° mismatch (down arrow). Placing one leg down has very little effect.

**Figure S5.**
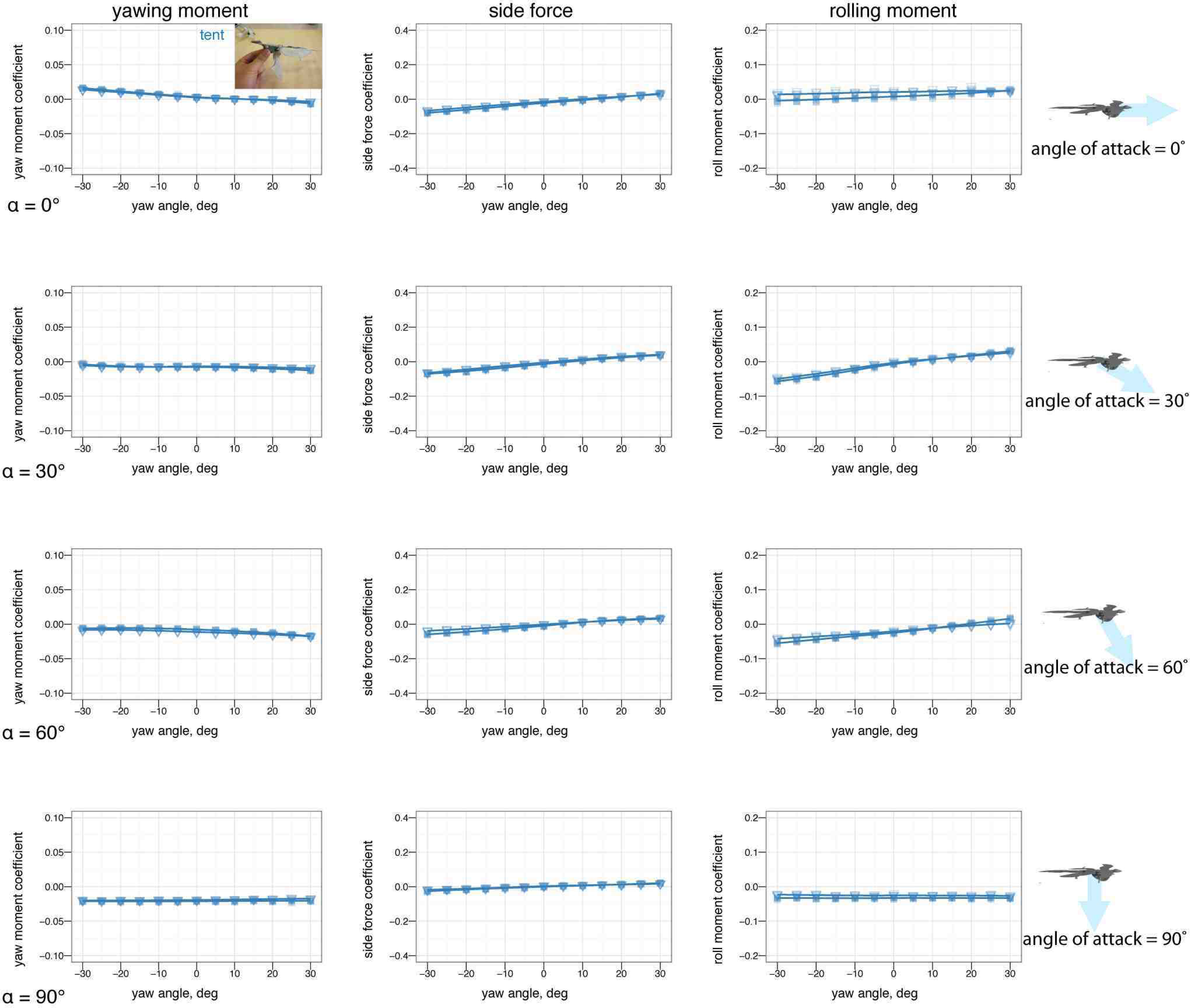
Asymmetric one leg down (leg *arabesque)* effect on yaw without leg or tail feathers. Baseline tent position (solid square) versus one leg at 90° mismatch (down arrow). Placing one leg down had little effect; with no leg or tail feathers there is no effect.

**Figure S6.**
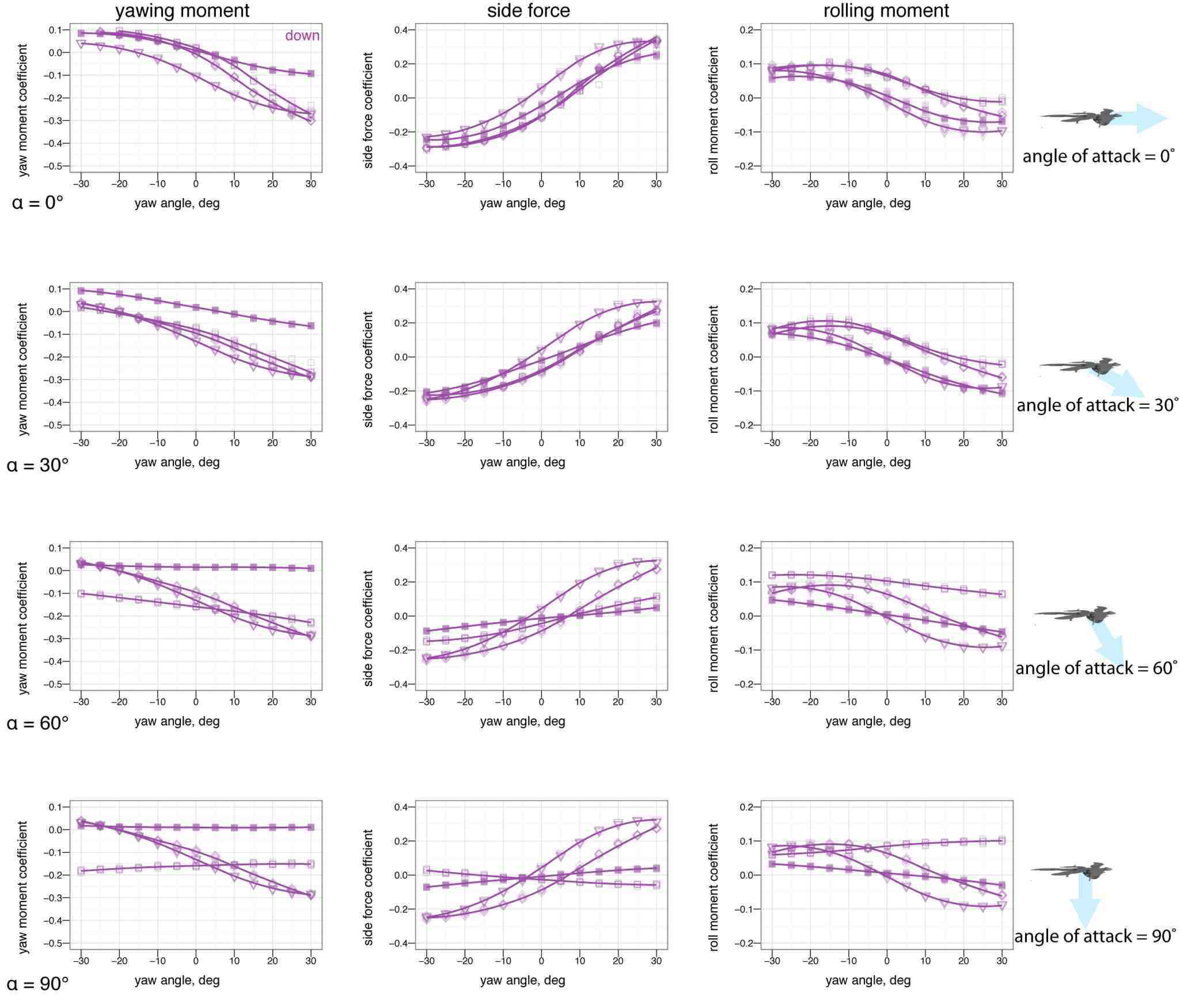
Asymmetric tail movement (lateral bending) effect on yaw, down posture. Baseline down position (solid square), tail 10° left (open square), tail 20° left (open triangle), tail 30° left (open diamond). The tail is effective at creating yawing moments.

